# Full 16S and 23S rRNA gene-based,strain-level resolution of the microbiota of a mock bacterial microbiome using ONT nanopore sequencing

**DOI:** 10.1101/2024.02.07.579414

**Authors:** Christopher J. Woodruff, Zhengming Zhang, Terence P. Speed

**Author notes:** Contributing authors.

## Abstract

**Purpose:** The feasibility of achieving near-strain taxonomic resolution and reliable microbiota strain-level abundance estimation using amplicon-based nanopore sequencing was investigated.

**Methods:** Denoising was applied to separate 16S and 23S genes extracted from a metagenomic dataset generated by Sereika et al. [9] using nanopore sequencing. Denoising used Kumar et al.’s [8] Robust Amplicon Denoising (RAD) and generated amplicon sequence variants (ASVs). The Sereika et al. data set was generated from the Zymo [10] D6322 7-bacterial species mock microbiome. A sub-sampling procedure generated additional datasets that allowed sensitivity assessment over multiple orders of relative abundance. Alignment to bespoke 16S and 23S rRNA gene databases, provided both identification and abundance information data for sub-species analyses.

**Results:** Sub-species identification was clearly achieved, both with the “even” D6322 dataset, and with the 4 sub-sampled datasets in which 3-orders of magnitude relative abundances were present. Multiple ASVs of length approximately 2500 bases were generated which gave perfect alignments to reference 23S rRNA genes. Similarly for the approximately 1500 base long 16S rRNA genes. Despite strain ambiguity for some species, the strain of each species known to be present was identified in all but 1 (of 70) cases examined for all species. A process to merge the 16S and 23S rRNA genes results reduced ambiguity, allowing better sub-species resolution, and better species, and sub-species, relative cellular abundance estimates.

**Conclusion:** Principled methods for dealing with strain-level ambiguity in microbiota analysis have been developed that are generally applicable. Their application to nanopore sequencing with the current state of Oxford Nanopore Technology, together with state-of-the-art denoising, allows near strain resolution of bacterial microbiota identity, and improved estimates of species, and sub-species, relative abundances.

## 1 Introduction

The early stages of microbiota analyses typically require identification of which microbial organisms are present and their relative abundance. This profiling is nowadays carried out either by shotgun sequencing of DNA from a sample of the microbiota (metagenomic sequencing, often accompanied by metagenomic assembly for identification), or by targeting of particular features common to the organisms present but which allow discrimination of those organisms. This latter approach is referred to as amplicon-based profiling, though it does not necessarily require any biochemical amplification process. Amplicon-based methods are efficient in their use of DNA and sequencing resources, but have traditionally lacked taxonomic resolution due to the limited length of the sequences being used. Ribosomal RNA (rRNA) genes have a particular advantage as there are multiple identical or almost identical such sequences in almost all bacteria, and hence an effective amplification process is achieved. Long read technologies, such as the Pacific Biosciences (PacBio) SMRT method - based on circular consensus sequencing (CCS) - and Oxford Nanopore Technologies (ONT) nanopore sequencing allow the use of sequences of such length that markedly better taxonomic resolution is feasible, as clearly demonstrated by De Oliveira Martins et al. [7] with respect to bacterial microbiota analysis. The high accuracy of recent PacBio HiFi sequencing has given strain resolution data on full 16S rRNA gene data. However, lower accuracy has limited the application of nanopore technology for sub-species resolution, apart from through methods that involve more complex preparation of DNA material prior to short read sequencing and then computationally derived synthetic long reads. Both Karst et al. [1] and Callahan et al. [2]) have demonstrated the value of such methods in giving mean error rates for nanopore sequencing of less than 0.005%. Such precision can clearly give sub-species taxonomic resolution, but, even with full length 16S and 23S rRNA gene alignments there still remains ambiguity in strain identification. The results from Karst et al. when applied to a real dataset - human faecal samples - were perhaps disappointing, in that with both 16S and 23S rRNA genes used strain resolution was still under 40%. Karst et al. report using the highest rank alignment to associate a reads-derived consensus sequence to a database strain. This is sound if there are no ties in this top ranking. The possibility of ties for both 16S and 23S genes simultaneously certainly exists and appears not to have been addressed.

Nanopore sequencing is far more available to those on low budgets, and is easily transportable. Extending its applicability to strain level microbiota resolution, without requiring more complex sample preparation methods, is likely to be globally beneficial to many researchers outside major cities, and those with highly constrained budgets. The results presented here demonstrate a valuable extension in taxonomic identification and associated abundance estimation for the bacterial component of microbiomes. In particular, the issue of multiple optimal alignments of any ASV to strains within the database has been comprehensively addressed. This reality of multiple optimal assignments at the strain level appears not to have been addressed previously.

### 1.1 Denoising of amplicon reads

Sequenced fragments result in a library of reads, the observed sequences of which are often slightly in error - that is, the reported, or observed, sequence is not exactly that of the section of the fragment from which it came. The errors in the reads constitute noise, and arise from both wet lab processes as well as the sequencing process (Quince et al. [3]). In microbiota and metagenome analysis “denoising” is a computational process applied to sequenced data from a microbiota sample that seeks to correct these errors from the wet lab and sequencing processes.

Denoising algorithm development has tended to combine general statistical methods of improving signal to noise with specific adaptations to the structure of the noisy processes. For instance, Quince et al. specifically address PCR reactions, including the formation of chimeras, as well as the modelling of errors arising from the sequencer technology - in their case, Ion Torrent technology. More recent work ([4], [5], [6]), focussed on Illumina NGS sequencing characteristics has improved on the modelling process and algorithms to produce code DADA2 [2]. Kumar etal. (2019)([8]) introduced RAD (Robust Amplicon Denoising) specifically focussed on the Pacific Biosciences RS II sequencer which had a weakness in dealing with homopolymers.

The error rate achievable from nanopore sequencing with the paired UMI method of [1] is of the order of 100 times lower than raw current error rates for such sequencing. Denoising using the RAD denoiser is shown here to return sequences that are generally at least the equivalent of about 10 times lower in error rate.

### 1.2 Amplicon Sequence Variants

Much microbiota analysis has used the concept of an Operational Taxonomic Unit (OTU) as an efficient representation of reads with similar sequences. OTUs are defined through an alignment and clustering process, coupled with a distance thresholding in the cluster space. More recently Amplicon Sequence Variants (ASVs) have been proposed ([4], [6]) as an alternative to OTUs as the basic analysis unit for sequence-based microbiota profiling that is directed towards phylogenetic classification. ASVs are explicit sequences, and if the data set from which they are derived is adequate they will be exactly the sequence of the genome from which they were derived. Being exact sequences of parts of a genome, any true ASV is independent of the data from which it is derived - though the set of proposed ASVs derived from any specific dataset clearly will depend on that dataset, in so far as they fail to be true ASVs. OTUs, however, are intrinsically dependent on the dataset from which they are derived. Phylogenetic trees can be derived in terms of ASVs, with the ASV effectively acting as a taxon, or node in the phylogenetic tree. Given a phylogenetic tree whose nodes are characterised in terms of ASVs, associating reads from some sample to locations on the tree requires only computing a distance measure to the ASVs characterising the tree, and noting the minimum such distance across the set of ASVs. In the remainder of this paper we use the term ASV, without further caveating, to refer to the output of a denoiser that attempts to derive ASVs.

In order to undertake amplicon-based bacterial phylogenetic analyses it is necessary to have amplicons from which data-independent phylogenetic tree nodes can be deduced. Generally, this cannot be achieved at the species level with short read sequencing as the subregions - V2, V3, etc. - of the 16S rRNA gene are often identical across species and even across genera (depending on the actual content of the microbiota being considered). However, if longer sequences are used, there is an increasing likelihood that species-level discrimination will be achieved - and some sub-species resolution ([1],[2]). These various long-read-based methods all rely on either rather complex synthetic long-read generation followed by NGS sequencing (e.g. LoopSeq, see [2]), or PacBio Sequel II or later sequencing. Such methods are not as generally available as sequencing based on the ONT nanopore technology. However, nanopore technology has not had sufficient acccuracy to provide the high taxonomic resolution of these other methods. Recently, it was shown ([9]) that the ONT R10.4 sequencing technology could provide metagenomic-assembled-genomes (MAGs) of a quality almost indistinguishable from that of the latest PacBio technology, and better than that based on NGS datasets. The following work is an investigation of whether the same quality of data may allow sub-species analyses of microbiota using full nanopore-produced, 16S and 23S rRNA gene-based sequences.

## 2 Methods

### 2.1 Extraction of 16S and 23S rRNA gene sequences from reference genomes

Primer sets targeted to low-entropy regions of the bacterial 16S and 23S rRNA genes were defined (Supplementary Material section 5). For the 16S rRNA gene subunit the complete genome was scanned with stepped, overlapping windows of window size 3000 bases and overlap 1600 bases (for the 23S rRNA gene: window size 4500, overlap 3000 bases). Each window was processed to identify whether it was likely that a 16S rRNA gene sequence lay within it. This involved checking for good quality alignment of at least one primer sequence from each of six primer sets, where a primer set targeted a single location (e.g. the V2 5’ location). If promising candidate boundary sets were obtained at this stage,multiple conditionals allowed reliable identification of all subunits within a genome. The sequences of all 16S rRNA genes identified were extracted, tagged according to species and indexed according to position from 5’ to 3’ on the reference genome. A completely analogous process was used for the 23S rRNA gene. While the process is largely unnecessary for 16S rRNA genes, it is necessary for 23S rRNA genes, as there is rather limited established reference data for 23S rRNA gene sequences. Comparison of 16S rRNA gene results against the rrnDB of Stoddard et al [12] showed perfect concordance, giving confidence in this novel search method within the 23S rRNA gene.

### 2.2 Construction of Reference Databases

For each species approximately 10 strains were selected randomly from Refseq complete genomes for bacteria. As there are 7 species in the mock microbiota this gives approximately 70 strains, with strains having between 2 and 14 16S rRNA gene or 23S rRNA gene sequences. The method of sub-section 2.1 is used to extract all 16S rRNA gene and 23S rRNA gene sequences.

The database for 16S rRNA gene studies is simply the set of all 16S rRNA gene sequences extracted from the strains selected. Computationally it would be more efficient to retain only a set of unique sequences from each strain, but this has not been done as this study has only used a small database and so gains in computational efficiency would be trivial. The 23S rRNA gene database is constructed in the same manner.

Note that the extractions of 16S and 23S rRNA genes are run independently, even though most 16S rRNA gene have a closely-located 23S rRNA gene This allows reliable identification of these genes in those bacterial species, such as Cereibacter sphaeroides, that do not have conventional ribosomal RNA (rrn) operons. It is a simple matter to additionally process the gene boundaries’ data to identify rrn operons, or to extract the Internally Transcribed Spacers (ITSs).

### 2.3 Datasets of nanopore-sequenced reads

Sereika et al. [9] generated metagenome datasets from a variety of different microbiomes. One was the Zymo D6322 mock microbiome which has equal masses of DNA from 7 bacterial species. This was sequenced with the ONT R10.4.1 pore on a Prometh*ION ^T^ ^M^* sequencer, giving approximately 3 million reads. A small proportion of these reads will have either a full 16S or 23S rRNA gene - and possibly both. Identification and extraction of these allows construction of 16S and 23S rRNA gene datasets which are synthetic amplicon datasets of the D6322 “even” bacterial mock microbiome.

Extraction of the rRNA genes from reads is closely related to the method of extracting the same rRNA genes from strains described in sub-section 2.1 However there is no stepped window - each read is processed in a single step. The same set of primers is used as for the reference database development, but less stringent tolerancing is used since reads are subject to a higher error rate than reference genomes. Also, for computational simplicity, no more than one 16S rRNA gene and one 23S rRNA gene is extracted from a read.

Our in-silico processing identified approximately 20,000 reads that had a full 16S rRNA gene sequence and a similar number had full 23S rRNA gene sequences (some had both). A library of such reads was constructed for each of these rRNA genes. RAD provided length and quality filtering of these reads which resulted in 4503 16S rRNA gene reads and 3707 23S rRNA gene reads being processed to generate ASVs. This set of filtered reads are referred to hereafter as the primary D6322 (16S rRNA gene or 23S rRNA gene) datasets. Length filtering involved lower bound setting (1200 for 16S rRNA gene and 2000 for 23S rRNA gene), while quality filtering - which is based on the Phred score returned by the basecaller - used the default value of an error rate of 1%. Since the 16S rRNA gene reads are approximately 1500 bases in length and the 23S rRNA gene are about 2400 bases the expected maximum numbers of errors is about 15 and 24, respectively.

As noted earlier the mock microbiome has proportions of the different species that are approximately the same, as there is equal DNA mass of each. To assess sensitivity of the method over orders of magnitude difference in relative abundance a sub-sampling process on the primary datasets was implemented. Knowing the quality of the ASVs (Section 3.1 and the distance between species (Table 11)), coupled with further detailed inspection of blastn alignment output, (see Supplementary Material Table 13 and Tables 14 to 19) we are confident that the species-level assignment of reads by RAD for the D6322full datasets is exact or very close to exact.

The sub-sampling involved randomly choosing a specified fraction of the reads for two or three species while retaining all reads from the other species. The sub-sampling factors used were (0.5, 0.2, 0.1, 0.02, 0.01). The process was not random as it was considered not likely that ASVs of reasonable quality would be returned if less than about 10 reads were available from a species. Also, if the Levenshtein distance between the strain’s 16S rRNA gene or 23S rRNA gene sequences was more than about 3 then multiple ASVs would tend to emerge from the denoising process. The set of downsampling values chosen is shown in Table 2.

**Table 1.** Reads assigned to each species from RAD denoising of 16S and 23S rRNA genes.

| ID | NumFilt | Bs | Ef | Ec | Lm | Pa | Se | Sa |
| --- | --- | --- | --- | --- | --- | --- | --- | --- |
| D6322full_16S rRNA gene | 4503 | 558 | 938 | 478 | 798 | 223 | 979 | 529 |
| D6322full_23S rRNA gene | 3707 | 379 | 739 | 454 | 505 | 192 | 833 | 605 |

Reads assigned to each species from RAD denoising of the D6322full 16S rRNA gene and 23S rRNA gene datasets. Species are Bs=*Bacillus subtilis*, Ef=*Enterococcus faecalis*, Ec=*Escherichia coli*, Lm=*Listeria monocytogenes*, Pa=*Pseudomonas aeruginosa*, Se=*Salmonella enterica*, Sa=*Staphylococcus aureus*.

**Table 2.** Sub-sampling weights to generate datasets sub1 - sub4.

| Dataset | Bs | Ef | Ec | Lm | Pa | Se | Sa |
| --- | --- | --- | --- | --- | --- | --- | --- |
| sub1 | 0.1 | 1 | 1 | 1 | 1 | 0.5 | 0.2 |
| sub2 | 0.5 | 0.01 | 0.1 | 1 | 1 | 1 | 1 |
| sub3 | 1 | 0.02 | 1 | 1 | 1 | 1 | 0.02 |
| sub4 | 1 | 1 | 0.1 | 1 | 1 | 1 | 0.02 |

Weights assigned to each species for subsampling of the D6322full 16S rRNA gene and 23S rRNA gene datasets. Species are Bs=*Bacillus subtilis*, Ef=*Enterococcus faecalis*, Ec=*Escherichia coli*, Lm=*Listeria monocytogenes*, Pa=*Pseudomonas aeruginosa*, Se=*Salmonella enterica*, Sa=*Staphylococcus aureus*.

Primary D6322 datasets that used only 16S and 23S sequences from reads that contained both were also generated. However the number of reads available was too small to provide good data for the sub-sampling undertaken. Such a limitation would normally not arise with a dataset generated specifically for amplicon-based microbiota analysis.

### 2.4 Denoising and ASV alignment

Sequences of the fragments with 16S rRNA gene and 23S rRNA gene are denoised using Kumar et al.’s Robust Amplicon Denoising (RAD) code. This process generates ASVs which typically are of a length close to, but slightly shorter, than the sequences in the reference database. These ASVs are then aligned against the relevant rRNA gene database, blastn being the code used. For each ASV this has returned approximately 50-100 sequences across our various datasets, one or more of which are exactly or very closely identical to actual sequences of the organisms present, based on exact alignments (most commonly) or almost always no more than 4 mis-matches or gaps (see Sub-Section 3.1). Sub-Section 3.1 presents data on how closely the ASVs determined using RAD align with reference genomes from which they are assumed to derive. For each ASV, RAD also returns a count of how many reads are best associated to it, and - optionally - the identity of each such read. This detailed information from the alignment process underpins the sub-sampling process described in Section 2.3 for producing datasets sub1 to sub4.

### 2.5 Identification and abundance estimation

Alignment of an ASV to the reference database returns a set of strain rRNA gene names and alignment metadata for each item returned. Note that, for any strain in the DB, each rRNA gene sequence of that strain is present. The strain, or strains, with the highest ranked alignment (including ties) is deemed to be possibly present. This process is followed for each ASV. Noting that each ASV has a count of reads associated with it, an initial count for each strain is generated. The count assigned at this stage involves making an assumption that the ASV only arises from a single strain, but if the same ASV is optimally aligned to multiple strains we cannot resolve which strain this should be based purely on this alignment data. Therefore each strain determined to be possibly present is recorded as present with the full count deriving from the ASVs that align optimally to it being assigned to any such strain. Note that the estimated count for the strains that are actually present must be such that the sum of those counts equals the full count. Thus the estimated strain count is dependent on decisions about which one or more of the possibly present strains are presumed to be actually present. We identify, therefore, two types of ambiguity - what we will call initial ambiguity which is the multiplicity of possibly present strains identified by optimally aligned ASVs, and residual ambiguity which is the multiplicity of possible groupings of strains following a decision process - soon to be described - applied to the possibly present information to desirably reduce the initial ambiguity. Section 2.6 describes the decision-making process applied when we merge information from the separate 16S rRNA gene analysis and the 23S rRNA gene analysis. This generates residual ambiguity which is shown to be often smaller than the residual ambiguity from un-merged data and also the initial ambiguity.

Although alignment will often apparently - and sometimes correctly - distinguish between rRNA genes within a strain we have not made use of such data in this work. This is further discussed in Section 4.

### 2.6 Principles for merging 16S and 23S rRNA gene strain level data

Merging will be shown in section 3.5 to reduce ambiguity of strains for some species. It also has implications for strain and species abundance estimation. Here we state the principles we have applied to develop the merging process adopted, detail a set of rules developed to implement those principles, and illustrate their operation with several examples.

It is assumed throughout this work that any optimal alignments of a particular ASV only occur for strains of a single species. This has been observed to be the case for the strains populating the databases used in this work, and is supported by the material of sub-section 3.2. A broader assessment of the validity of this assumption has not been undertaken. Real data for such a broader assessment is not currently available. Some indication of the likely validity could be achieved using 16S rRNA gene reference databases and examining edit distances between 16S rRNA gene sequences from different species.

The ASVs considered in this work are close to the length of the full 16S rRNA gene or 23S rRNA gene amplicons. If a strain is present in the sample to the extent that hundreds of reads are sequenced from this strain then, typically, multiple ASVs will be generated. As a consequence there are often multiple ASVs that optimally align to that strain. If two strains are very similar and are both present in the microbiota, the ASVs derived from them could be such that reads from one actually best align with ASVs from the other. Thus we may not only have multiple ASVs aligning to a strain, but we may have some ASVs that have the same pattern of optimal alignment to a set of strains. Note that the quality of the optimal alignment may be different for different ASVs.

#### 2.6.1 Merging Principles and Derived Rules

Two principles are proposed, followed by a set of rules to implement those principles. The term “possibly present” is used to describe any strain having at least one 16S rRNA gene or 23S rRNA gene ASV optimally aligning to it.

Principle 1: If any strain is found to be possibly present both from 16S rRNA gene and 23S rRNA gene alignment of ASVs to the reference database, it is more likely to be present than is a strain found possibly present only by one of the 16S rRNA gene and 23S rRNA gene alignment processes. This is the basic principle underlying the ambiguity reduction sought by merging. We label a strain that is found possibly present by both 16S rRNA gene and 23S rRNA gene alignments as a “shared strain”, and consider a shared strain to more likely be present than a strain which is not shared.

Principle 2: In the absence of information for discrimination between strains choose the minimal number of strains consistent with the data.

These principles have been implemented using the following rules. Further refinement of the handling of both shared and not-shared strains is needed for abundance quantification, and this appears in the rules.

Rule 1: Any shared strain, or strains, of a species, are preferentially treated in assignment of reads from ASV sets in abundance calculations. Conseqently, if an ASV set is associated with at least one shared strain it is considered not to be associated with any non-shared strain.

Rule 2: Choose the minimal number of shared strains to account for the maximal number of ASV sets.

Rule 3: If two or more shared strains from a minimal set of strains consistent with Rule 2 themselves have an ASV that is optimally aligned to each of them, then the read count associated with such an ASV is partitioned across these strains, such that the cellular abundances of such strains are equal.

Rule 4: Any ASV that is associated with a shared strain, or strains, has all of its associated reads attributed only to the shared strain or strains.

Rule 5: An ASV set that is not associated with a shared strain has its read count assigned to exactly one of those strains to which it is optimally aligned. Ambiguity of such strains is resolved by a random choice from those still-ambiguous strains.

Two alternate treatments of not-shared strains have been considered. One is to report all such strains, with an abundance based on equal apportioning of ASV set counts. The other reports an upper and lower bound on cellular abundance through selective retention of strains that maximise or minimise this quantity.

#### 2.6.2 Illustrations of Application of the Merging Rules

Illustrations of the application of these rules is provided in the following. Consider binary matrices, B, for a species where the columns of B correspond to ASV sets associated with the species, and the rows correspond to strains. Note that Cases 1 and 2 are modified representations of the true data so as to allow better illustration of the rule application process.

##### Case 1: *Escherichia coli*

Table 3 shows such matrices for the *Escherichia coli* 16S rRNA gene and 23S rRNA gene data. Note that indexing of ASV sets for the 16S rRNA gene is independent of that for the 23S rRNA gene.

**Table 3.** ASVsets presence matrices for strains - *Escherichia coli* example.

| ASV set index | 10 | 11 | 12 | 14 | 15 | 16 |
| --- | --- | --- | --- | --- | --- | --- |
| 000913 | 1 | 0 | 1 | 1 | 0 | 1 |
| CP062729 | 0 | 1 | 0 | 0 | 0 | 0 |
| CP024141 | 0 | 1 | 0 | 0 | 0 | 0 |
| CP048821 | 0 | 0 | 0 | 0 | 1 | 0 |
| CP029747 | 0 | 0 | 0 | 0 | 1 | 0 |

| ASV set index | 10 | 11 | 12 | 13 |
| --- | --- | --- | --- | --- |
| 000913 | 0 | 0 | 0 | 1 |
| CP012759 | 1 | 0 | 1 | 0 |
| CP048821 | 0 | 1 | 0 | 0 |

Presence/absence matrices of ASV sets optimally aligning to *Escherichia coli* strains for the 16S rRNA gene (upper) and 23S rRNA gene (lower). Note that indexing for the 16S data is unrelated to that for the 23S data.

It can be seen that the first and fourth strains (000913 and CP048821, respectively) in the 16S rRNA gene table are identical to the first and third strains in the 23S rRNA gene table Therefore, by Rule 1, both strains are considered shared. Next we note that, for the 16S rRNA gene amplicon, strain 000913 has ASV sets 10, 12, 14, and 16 associated with it and these are not associated with the other shared strain. Therefore strain 000913 has all reads assigned to those ASV sets considered to arise from this strain. The other shared strain, CP048821, has ASV set 15 only associated with it.

That ASV set is also associated with strain CP029747. However, strain CP029747 presence is not supported by the 23S rRNA gene data, and hence, by Rule 4, those reads associated with ASV set 15 are all assigned to the shared strain CP048821. Only ASV set 11 remains unaccounted for. We have no data to differentiate between strains CP062729 and CP024141. Rule 5 would allow either, but not both, of these strains. Rule 5 provides a resolution path in this case - random choice between 2 strains. Thus the decision making process gives two groups of strains that could meet the requirements. The residual ambiguity for 16S data is therefore 2 under merging, while there is only a single combination of strains that is valid for the 23S data, so resulting in a residual ambiguity of 1 for the 23S data. Note that, in the absence of merging, applying Rule 5 to the 16S data would result in 4 possible triplets of strains meeting that Rule, and hence the residual ambiguity without merging is 4 for the 16S data, whereas the initial ambiguity was 5. In this case merging has decreased the residual ambiguity from 16S data for this species from 4 to 2.

Application of these rules implements a decision process giving one or more minimal sets of strains. The residual ambiguity is the number of such minimal sets. Each such set results in a modified B matrix we will call the R matrix. The R matrix will have non-integer entries in some cells when shared strains have an ASV set in common. In this *Escherichia coli* illustrative case that is not the case. Table 4 shows one such resultant R16 matrix.

**Table 4.** 16S ASV set abundance allocation proportions for ASV sets.

| ASV set index | 10 | 11 | 12 | 14 | 15 | 16 |
| --- | --- | --- | --- | --- | --- | --- |
| 000913 | 1 | 0 | 1 | 1 | 0 | 1 |
| CP062729 | 0 | 1 | 0 | 0 | 0 | 0 |
| CP024141 | 0 | 0 | 0 | 0 | 0 | 0 |
| CP048821 | 0 | 0 | 0 | 0 | 1 | 0 |
| CP029747 | 0 | 0 | 0 | 0 | 0 | 0 |

One case of 16S ASV set abundance allocation proportions for ASV sets optimally aligning to *Escherichia coli* strains for the 16S rRNA gene subunit. Note that, by Rule 4, either CP062729 or CP024141 must be assigned 1, and the other 0 - the choice is random, and CP062729 was selected here.

##### Case 2:*Staphylococcus aureus*

A second example is provided by S.aureus. Table 5 gives both the 16S rRNA gene and 23S rRNA gene B matrices, together with the number of operons of each strain. There are 6 strains identified from the 16S rRNA gene analysis, and 8 by the 23S rRNA gene analysis. All 6 of the 16S rRNA gene strains are also amongst the 8 23S rRNA gene strains. Thus there are 6 shared strains.

**Table 5.**
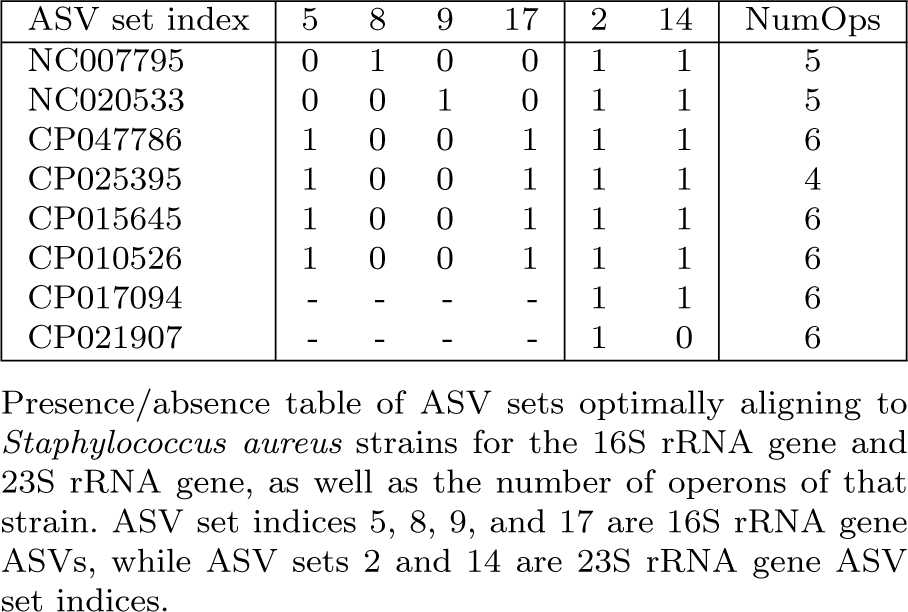
ASVsets presence matrices for strains - *Staphylococcus aureus*.

For the 16S rRNA gene strains 1 and 2 (NC007795, NC020533) each have a single ASV set associated, and that set is unique to the particular strain. However strains 3 to 6 each share two ASV sets. By rule 2 any one of these can be chosen, providing 4 possible 3-strain solution sets. There are no shared ASV sets within any of these solution sets. The 23S data allows no discrimination between strains. Hence the 4 solution sets from the 16S data would also apply for the 23S data. This results in shared 23S ASV sets for each of these solution sets and hence apportioning of the 23S ASV counts. Since the operon count is different for different strains the strain cellular abundance estimates vary according to which one of strains CP047786, CP025395, CP015645 and CP010526 are in the solution set being considered-specifically whether it is the 4-operon CP025395 or not. By rule 3 the 23S ASV sets 2 and 14 have their counts distributed with weightings (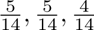)) if the strain is CP025395 otherwise (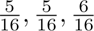). Sub-section 4.1 of the Supplementary Material formalises this apportioning process.

Under the merging process the residual ambiguity for the 16S data here is 4 as only shared strains are necessary to cover all relevant ASV sets, and there are 4 triplets of strains that provide full accounting of all ASVs - both 16S and 23S - optimally aligning to these strains. Without merging there are 4 possible solutions meeting Rule 4 for the 16S and 7 for the 23S. Thus, without merging, the initial ambiguity for 16S data is 4, the residual ambiguity 4 and for 23S the initial ambiguity is 8 and residual ambiguity 7.

The resultant R16 and R23 matrices for the case where the strain CP025395 is included are given in Table 6.

**Table 6.**
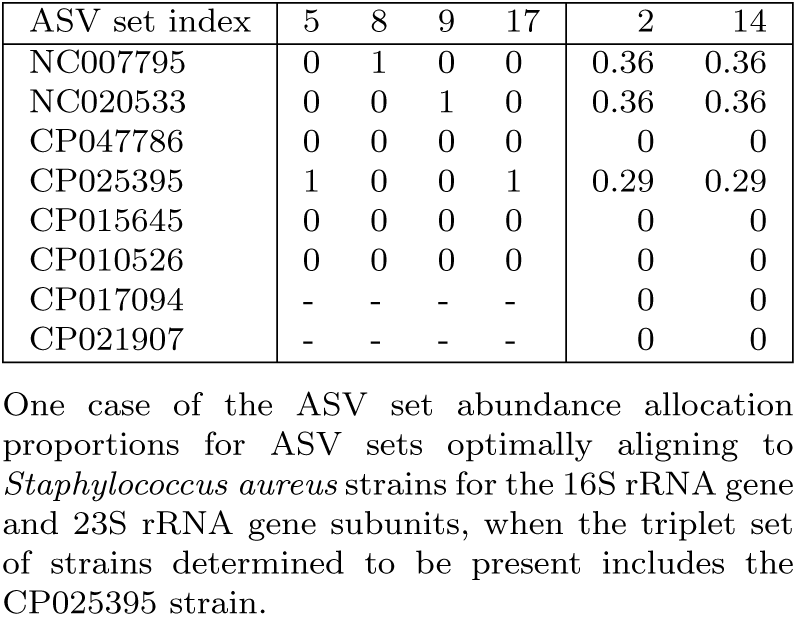
ASV set abundance allocation proportions for Case 2 strains.

For the 23S rRNA gene strains 7 and 8 (CP017094, CP021907) are not shared, and have no ASV sets not covered by the shared strains, hence they are not retained. The strain raw counts are determined as the weighted sum of the ASV sets counts, with weights given by the row of the R matrix corresponding to that strain. Cellular abundances are then calculated by dividing by the respective number of operons. This example highlights how species cellular abundances estimates are dependent on what strains are present when different strains have different numbers of rRNA genes.

##### Case 3: *Enterococcus faecalis*

This case is exactly the merging situation for the primary D6322 16S and 23S datasets, and provides an explanation for the cellular abundance values in Table 14 for *Enterococcus faecalis*. The 16S rRNA gene dataset gave 10 strains all of which had the same 2 ASVs. The 23S rRNA gene dataset gave 7 strains some with 1 and some with 2 ASVsets. Table 7 shows the B16 and B23 matrices. The number of operons is 4 for all strains.

**Table 7.** ASVsets presence matrices for strains - *Enterococcus faecalis*.

| ASV set index | 1 | 21 | 1 | 4 | 5 |
| --- | --- | --- | --- | --- | --- |
| D6322 | 1 | 1 | 1 | 0 | 1 |
| CP008816 | 1 | 1 | 1 | 0 | 0 |
| CP060796 | 1 | 1 | 0 | 0 | 1 |
| 017312p | 1 | 1 | 0 | 1 | 1 |
| 004668 | 1 | 1 | 0 | 1 | 1 |
| CP098743 | 1 | 1 | 0 | 0 | 1 |
| CP110074 | 1 | 1 | 0 | 1 | 0 |
| CP110059 | 1 | 1 | - | - | - |
| CP091904 | 1 | 1 | - | - | - |
| 017316 | 1 | 1 | - | - | - |

ASV set abundance allocation proportions for ASV sets optimally aligning to *Enterococcus faecalis* strains for the 16S rRNA gene and 23S rRNA gene subunits

There are 7 strains shared. For the 16S data there are 3 strains not shared, and zero for the 23S data. The merged decision process generates 3 doublet set-of-strains solutions - (D6322, CP110074), (CP008816, 0173120), and (CP008816, 004668) - each having no overlap in 23S ASV sets, but complete overlap in 16S ASV sets. For both 16S and 23S data, after merging, the residual ambiguity is therefore 3. Without merging the 16S data has residual ambiguity equal to the initial ambiguity of 10, while the 23S data has a initial ambiguity of 7 but residual ambiguity of 3.

The read count associated with the 16S rRNA gene ASV set 1 is 915, ASV set 21 is 23, while for the 23S rRNA gene ASVsets 1,4 and 5 the counts are 376, 207 and 156 respectively. Consider the merged solution of doublet (D6322, 017312). Under the merging process the 16S ASV counts are equally divided between two strains as these have the same number of operons. There is no across-strain overlap of ASV sets for any of the 23S ASV sets. All strains have four operons, so - for instance-the cellular abundance of the D6322 16S rRNA gene is (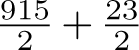)*/*4 = 117.5. Table 8, summarises the strain-level results for the doublet of strains (D6322, 017312).

**Table 8.**
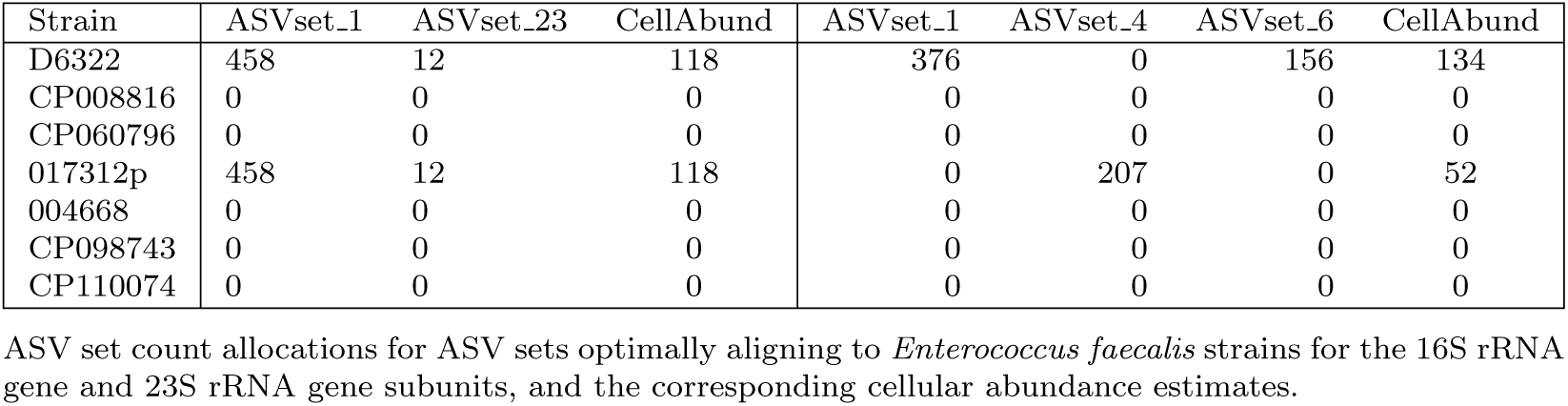
Strain count allocation and cellular abundance estimates for one doublet of strains.

| Strain | ASVset_1 | ASVset_23 | CellAbund | ASVset_1 | ASVset_4 | ASVset_6 | CellAbund |
| --- | --- | --- | --- | --- | --- | --- | --- |
| D6322 | 458 | 12 | 118 | 376 | 0 | 156 | 134 |
| CP008816 | 0 | 0 | 0 | 0 | 0 | 0 | 0 |
| CP060796 | 0 | 0 | 0 | 0 | 0 | 0 | 0 |
| 017312p | 458 | 12 | 118 | 0 | 207 | 0 | 52 |
| 004668 | 0 | 0 | 0 | 0 | 0 | 0 | 0 |
| CP098743 | 0 | 0 | 0 | 0 | 0 | 0 | 0 |
| CP110074 | 0 | 0 | 0 | 0 | 0 | 0 | 0 |

ASV set count allocations for ASV sets optimally aligning to *Enterococcus faecalis* strains for the 16S rRNA gene and 23S rRNA gene subunits, and the corresponding cellular abundance estimates.

The species cellular abundance for *Enterococcus faecalis* is simply the sum of the strain cellular abundances - 235 from 16S data and 186 from 23S data. Because all strains have the same number of operons the species abundance does not, in fact, depend on the distribution across strains. This example, however, demonstrates the application of the rules implemented.

## 3 Results

The datasets available for analysis are partially characterised in Table 9. The species counts entered for the sub-sampled datasets assume that the results from species-level identification and observed abundances of the primary D6322 datasets are exact in both respects. The counts for the species of the sub-sampled datasets provide the basis for determining an expected set of species proportions for the dataset. The expected proportions for the primary D6322 are computed based on the equality of DNA mass for each species in the original Zymo D6322 mock microbiome sequenced by Sereika et al.. In both cases the assumptions underlying these determinations will not be fully met.

**Table 9.** Read counts for datasets.

| Dataset | NumReads | NumFilt | Bs | Ef | Ec | Lm | Pa | Se | Sa |
| --- | --- | --- | --- | --- | --- | --- | --- | --- | --- |
| D6322full_16S rRNA gene | 15097 | 4505 |  |  |  |  |  |  |  |
| D6322full_23S rRNA gene | 16361 | 3607 |  |  |  |  |  |  |  |
| sub1_16S rRNA gene | 3089 | 3089 | 56 | 938 | 478 | 798 | 223 | 490 | 106 |
| sub1_23S rRNA gene | 2914 | 2914 | 38 | 739 | 454 | 505 | 192 | 416 | 101 |
| sub2_16S rRNA gene | 3295 | 3295 | 279 | 9 | 478 | 798 | 223 | 979 | 529 |
| sub2_23S rRNA gene | 2786 | 2786 | 190 | 7 | 454 | 505 | 192 | 833 | 505 |
| sub3_16S rRNA gene | 3066 | 3066 | 558 | 19 | 478 | 798 | 223 | 979 | 11 |
| sub3_23S rRNA gene | 2390 | 2390 | 379 | 15 | 454 | 505 | 192 | 833 | 11 |
| sub4_16S rRNA gene | 3555 | 3555 | 558 | 938 | 48 | 798 | 223 | 979 | 11 |
| sub4_23S rRNA gene | 2705 | 2705 | 379 | 739 | 45 | 505 | 192 | 833 | 11 |

Pre- and Post-filtering number of reads of D6322 in-silico Amplicon Datasets, together with the sub-sampled count of each species. Note that sub-sampling was applied to the RAD-filtered D6322full reads. The primary D6322 rows do not have counts entered as these are actually not known. The maximum value in each species column - separately for the 16S rRNA gene and 23S rRNA gene entries - is the result from estimating the counts using the methods described in this paper. Species are Bs=*Bacillus subtilis*, Ef=*Enterococcus faecalis*, Ec=*Escherichia coli*, Lm=*Listeria monocytogenes*, Pa=*Pseudomonas aeruginosa*, Se=*Salmonella enterica*, Sa=*Staphylococcus aureus*.

These 5 datasets per rRNA gene formed the basis for assessing how well this quality of long read data could provide taxonomic resolution and sample cellular abundance proportions in the mock microbiome. Table 9 summarises the datasets. Note that down-sampling was based on the species-level classification and count of reads from processing the primary 16S rRNA gene and 23S rRNA gene datasets.

A critical question is whether RAD is able to form useful ASVs for strains constituting a small proportion of the total reads. Table 10 presents the number of ASVs for each species for both the full 16S rRNA gene and 23S rRNA gene datasets and the various sub-sampled datasets. Clearly such ASVs can be generated. Not surprisingly the number of ASVs decreases as the number of sequenced reads decreases. Specifically, Spearman rank-order correlations between the number of ASVs generated and the subsampling rate were calculated for each of those species that were sub-sampled in multiple datasets - *Bacillus subtilis, Enterococcus faecalis*, *Escherichia coli* and *Staphylococcus aureus* - giving values of 0.72, 0.85, 0.86, and 0.96 respectively, all with p-values less than 0.02.

**Table 10.** Number of ASVs generated by RAD for each dataset.

| ID | P.aer. | S.aureus | E.coli | S.enteri. | E.faec. | L.mono. | B.subt. |
| --- | --- | --- | --- | --- | --- | --- | --- |
| D6322full.16S rRNA gene | 2 | 11 | 15 | 27 | 7 | 12 | 10 |
| D6322full.23S rRNA gene | 3 | 10 | 14 | 27 | 8 | 11 | 14 |
| sub1.16S rRNA gene | 5 | 3 | 18 | 14 | 13 | 6 | 3 |
| sub1.23S rRNA gene | 2 | 4 | 14 | 13 | 9 | 8 | 2 |
| sub2.16S rRNA gene | 3 | 14 | 3 | 33 | 1 | 11 | 7 |
| sub2.23S rRNA gene | 3 | 11 | 2 | 20 | 0 | 6 | 7 |
| sub3.16S rRNA gene | 3 | 1 | 15 | 28 | 1 | 9 | 6 |
| sub3.23S rRNA gene | 2 | 1 | 10 | 22 | 1 | 5 | 11 |
| sub4.16S rRNA gene | 5 | 1 | 3 | 26 | 20 | 4 | 7 |
| sub4.23S rRNA gene | 2 | 2 | 3 | 20 | 14 | 6 | 9 |

**Table 11.**
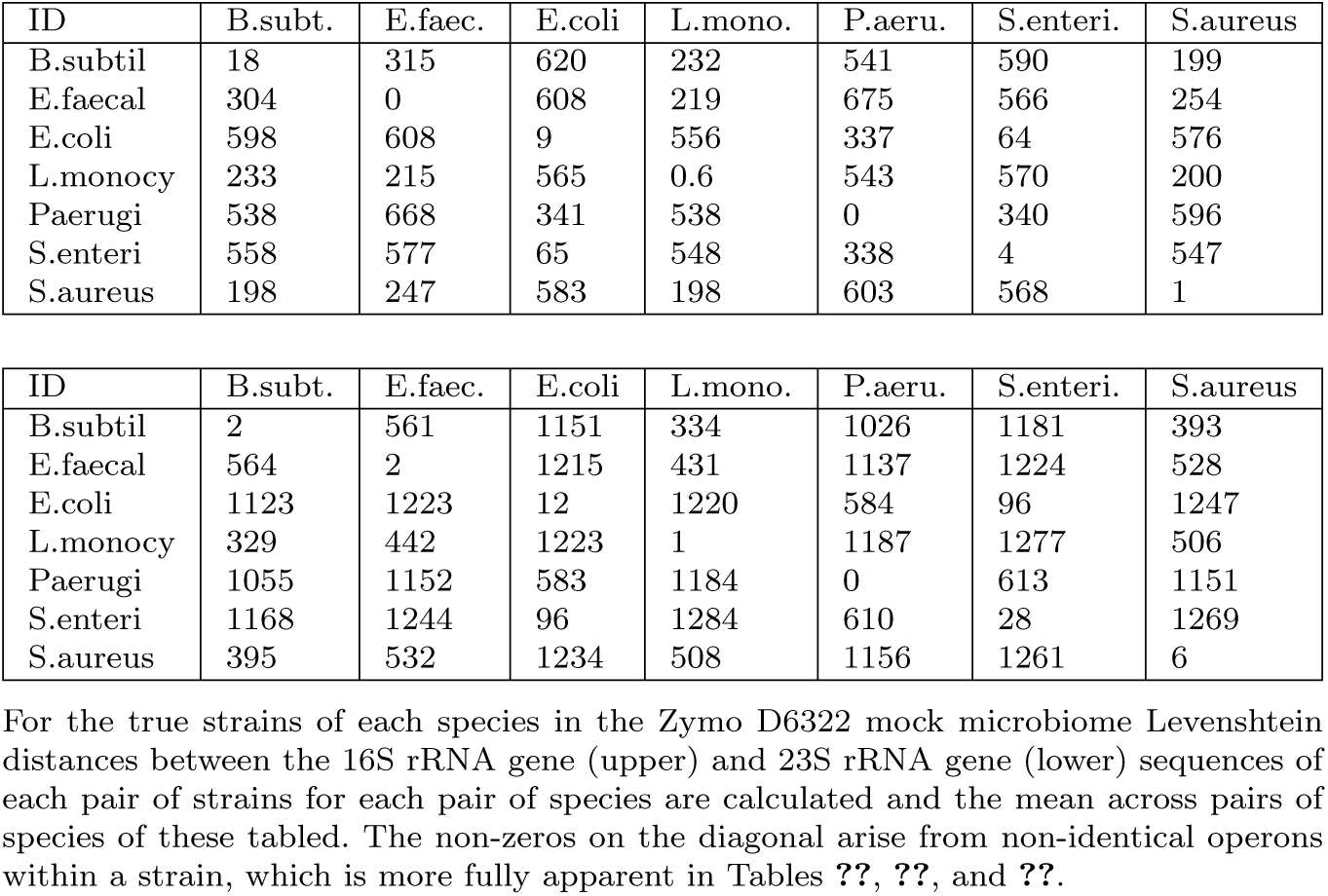
Mean Levenshtein distances between strains of each pair of species.

It is noted that, even in the cases where the reads of a species remain invariant between datasets, the number of ASVs for a particular species and subunit varies for the datasets - see *Listeria monocytogenes* and *Pseudomonas aeruginosa*, for example. It is presumed that this is a consequence of the stochastic nature of the ASV generation process in RAD.

Referring to Tables 9 (for read counts) and 10 (for ASV counts) it can be seen that species present at as small as 0.3% (*Enterococcus faecalis*, sub3, 16S) have at least one ASV generated, and this occurs reliably over several species having fewer than 1% of the number of reads in the dataset. It is also noted that the one case of a down-sampling factor of 0.01 resulted in no 23S ASV being generated that aligned to that species, though one ASV was generated for the corresponding 16S ASV. Thus RAD processing of 16S rRNA gene and 23S rRNA gene amplicons derived from nanopore-sequenced D6322 mock microbiota bacteria is effective in extracting meaningful ASVs with as few as about 10 reads present - see Table 10 - in a dataset of 3000 reads.

### 3.1 Accuracy of ASVs

The ability to resolve strains within a species depends on both the accuracy of the denoising process and the difference between the genomic sequences for the strains stored in the database. Here we present data on the former aspect, using counts of alignment errors - mismatches and gaps. The denoising process aims to produce exact sequences of the fragments that were sequenced. Our best guide to that is whether the alignment of an ASV to a database entry that is known to be that of a strain present in the mock microbiota gives a perfect alignment over a length close to the length of the database entry. Figure 1 presents summary data on the optimal alignments for each of the 16S rRNA gene ASVs derived for the primary D6322 dataset.

**Fig. 1.**
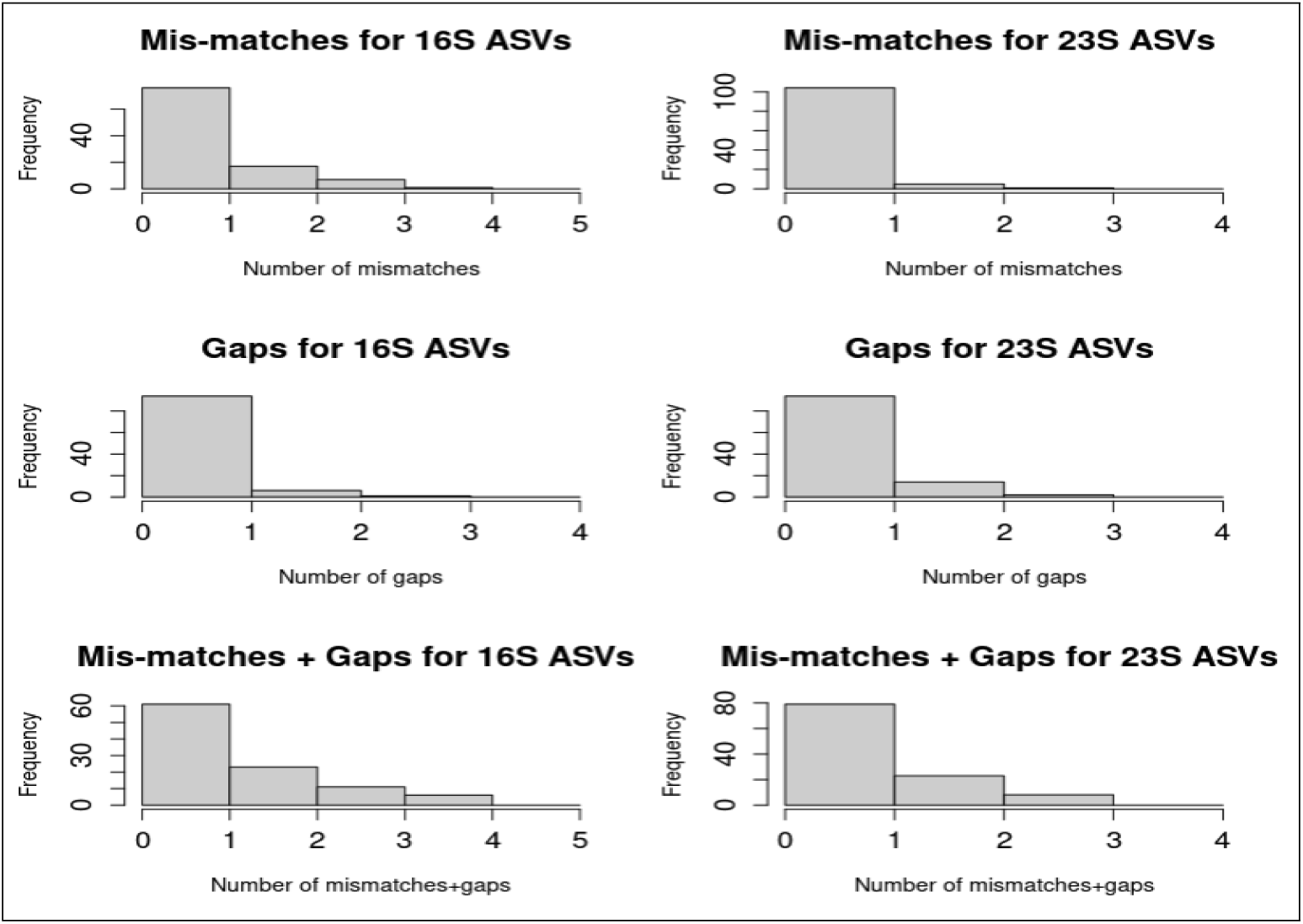
Histograms of mismatches and gaps for optimal alignments of ASVs against the reference databases for 16S rRNA gene and 23S rRNA gene amplicons of strains. There are 101 16S rRNA gene ASVs and 110 23S rRNA gene ASVs.

The majority of reads are associated with ASVs having zero gaps and zero mismatches - 57% for the 16S rRNA gene, and 61% for the 23S rRNA gene. (Supplementary Table 13). Even if there were no correlation between optimal 16S and 23S rRNA gene alignments this requires that more than 25% of reads would be associated with pairs of 16S and 23S rRNA ASVs that simultaneously align perfectly with strains. Under a common error rate for each location along such ASVs, and a Poisson model of errors, the error rate must therefore be less than 0.035%. Also, for both rRNA genes, fewer than 2% of reads are associated with ASVs having more than 3 gaps plus mismatches in their optimal alignments. More than 80% of reads are associated with ASVs differing by no more than 1 gap or 1 mismatch. Interestingly, the 23S rRNA gene ASVs are actually appearing to be of better quality than the 16S rRNA gene ASVs. Further detail on the matching of reference sequences and ASVs is presented in the Supplementary Material, Section 3.

### 3.2 Characteristics of the Reference Databases

Here we present data on the difference between the genomic sequences for the strains stored in the database, using the Levenshtein distance as the metric of pairwise distance between sequences. Evaluation of the inter-strain distances is useful only in the context of knowledge of the sort of distances that bracket different discrimination decisions regarding which strains an ASV aligns to well. Anticipating results of the following sub-section, we focus on Levenshtein distances both between 16S rRNA gene sequences and between 23S rRNA gene sequences that are less than 7.

Consider firstly the inter-strain distances for pairs of strains from different species. How different is the true strain of a particular species from the true strain of each other species in the mock microbiota? We compute the distance between each 16S rRNA gene (or 23S rRNA gene) sequence of the true strain of one species and the 16S rRNA gene sequences of the true strain of another species and then calculate the mean, standard deviation and range of this matrix of inter-species, inter-strain distances. Table 11 presents within and between species means for each of the 16S rRNA gene and the 23S rRNA gene. These results show how distant the species of this mock microbiome are. This large separation underpins the assumption that the species assignments of the D6322full dataset is highly accurate and so sub-sampling at the species level based on that is valid.

Now consider how the strains whose sequences are present in the database differ from the sequence of the strains, which we call the D6322 strains, that are actually in the microbiome Note that most bacteria have multiple ribosomal RNA operons (and hence 16S rRNA gene and 23S rRNA genes), and these generally are not all identical. Similarity, therefore, has been assessed for a single species by examining the Levenshtein distances between the set of 16S rRNA gene (or 23S rRNA gene) sequences of the D6322 strain and that of other strains of that species in the database. Of interest are cases where the distance is close to zero. Figure 2 provides a set of histograms showing the distribution of small distances between 16S rRNA gene sequences of the *Bacillus subtilis* strains from the D6322 strain. Strains that are more than a distance of 3 from an ASV for 16S rRNA gene sequences have been found to rarely give optimal alignments to that ASV. Thus, although not definitive, it is expected that only (non-D6322) strains having sequences with a distance from the D6322 strain of less than about 3 would possibly be identified (wrongly) as also present in the microbiota.

**Fig. 2.**
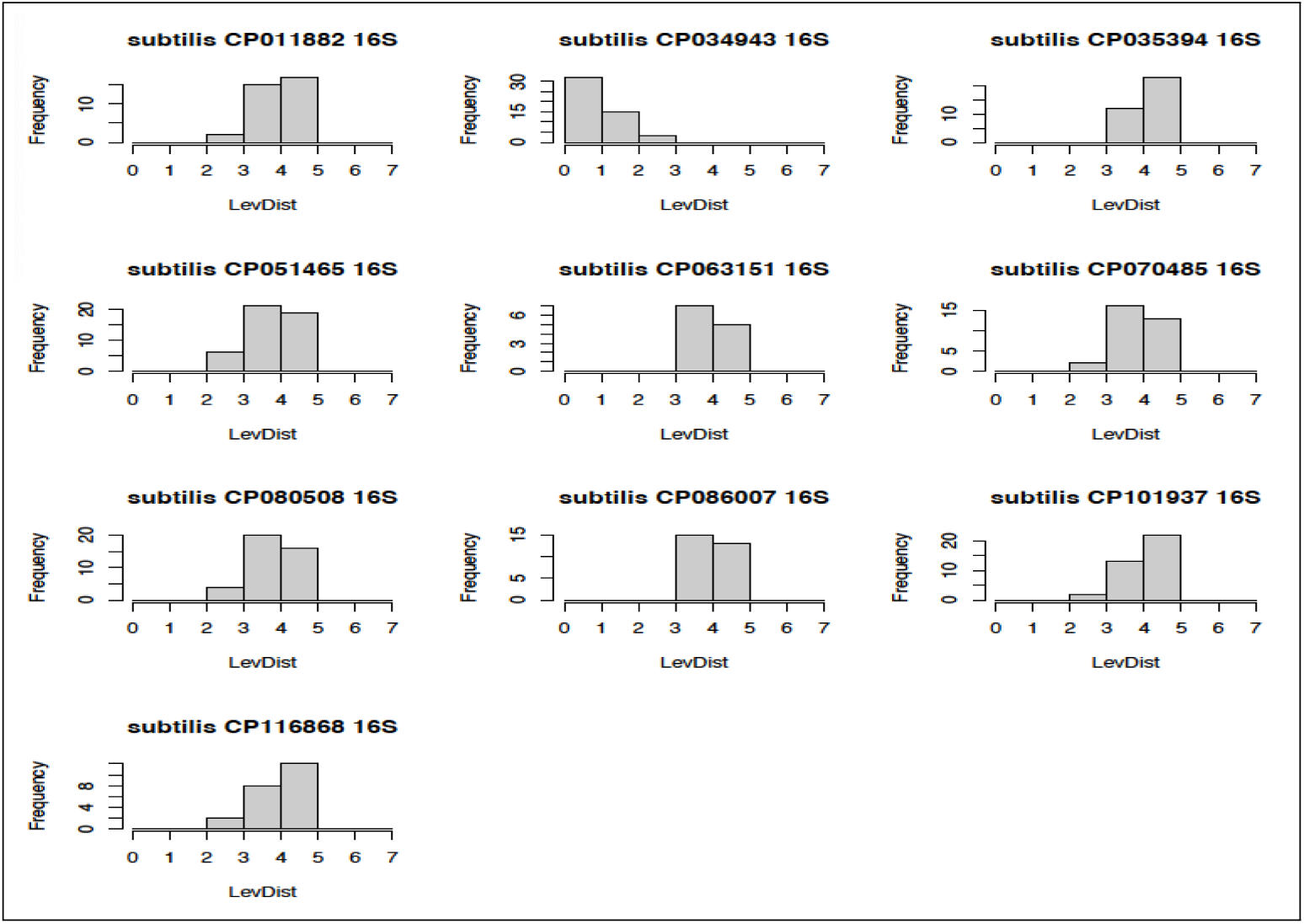
Levenshtein distance between 16S rRNA gene sequences of the D6322 B.subtilis strain and those of all other B.subtilis strains in the reference database. Only distances less than 7 are shown. Note that a bar from n to n+1 represents a count for distance n.

Finally, consider the implications of the reference database not having an entry corresponding to the species or strain from which some of the reads derived. The effect of removing all *Bacillus subtilis* strains - and hence the species - from the reference database was briefly investigated. Assignment of ASVs to the reference DB if the alignment percent identity (PID) was less than a threshold of 95% was implemented, with such ASVs being assigned to strain and species “other” and “Other”, respectively. The result, not surprisingly, was that all the reads previously assigned to strains of *Bacillus subtilis* were assigned to “other”. Hence no species proportions were altered, except for *Bacillus subtilis*. If no highly similar strain is present in the database it is still likely that reads from the strain would form at least one ASV (and probably multiple ASVs), and such an ASV, or set of ASVs, would have an optimal alignment probably to a strain within the correct species.

### 3.3 Strain-Level Identification

Separate, essentially identical analyses of the 16S rRNA gene and 23S rRNA gene datasets described in section 2.5 were undertaken. The optimal alignments of ASVs identify particular strains as possibly being present in the mock microbiota. For each ASV the strains (but not the operons) for which the optimal PID arose were retained. Table 12 presents, in columns 1 and 2, results from this stage of the identification process based on 16S rRNA gene data, while Table 13 give 23S rRNA gene data results.

**Table 12.** Strains and Abundance-related data from 16S rRNA gene ASVs of the primary D6322 dataset.

| Species | Strain | Raw Counts | Num. Ops | CellAbund |
| --- | --- | --- | --- | --- |
| *Bacillus subtilis* | D6322 NRRL B-354 | 558 | 10 | 56 |
| *Enterococcus faecalis* | NZ-CP110074-BE5 | 938 | 4 | 234 |
| *Enterococcus faecalis* | NZ-CP110059 BE16 | 938 | 4 | 234 |
| *Enterococcus faecalis* | NZ-CP098743 M9 | 938 | 4 | 234 |
| *Enterococcus faecalis* | NZ-CP091904 43-2 | 938 | 4 | 234 |
| *Enterococcus faecalis* | NZ-CP060796 EF-2001 | 938 | 4 | 234 |
| *Enterococcus faecalis* | NZ-CP008816 ATCC-29212 | 938 | 4 | 234 |
| *Enterococcus faecalis* | NC-017316.1 OG1RF | 938 | 4 | 234 |
| *Enterococcus faecalis* | NC-017312.1 62 | 938 | 4 | 234 |
| *Enterococcus faecalis* | NC-004668.1 V583 | 938 | 4 | 234 |
| *Enterococcus faecalis* | D6322 NRRL B-537 | 938 | 4 | 234 |
| *Escherichia coli* | D6322 NRRL B-1109 | 198 | 7 | 28 |
| *Escherichia coli* | NC-000913.3 K-12 | 280 | 7 | 40 |
| *Listeria monocytogenes* | D6322 NRRL B-33116 | 798 | 6 | 133 |
| *Pseudomonas aeruginosa* | D6322 NRRL B-3509 | 223 | 4 | 56 |
| *Pseudomonas aeruginosa* | NZ-LR134309 NCTC1290 1 | 223 | 4 | 56 |
| *Pseudomonas aeruginosa* | NZ-CP109931 PALA42 | 223 | 4 | 56 |
| *Pseudomonas aeruginosa* | NZ-CP054792 SE5431 | 223 | 4 | 56 |
| *Pseudomonas aeruginosa* | NZ-CP054791 SE5430 | 223 | 4 | 56 |
| *Pseudomonas aeruginosa* | NZ-CP030351 AR-460 | 223 | 4 | 56 |
| *Pseudomonas aeruginosa* | NZ-CP021774 Pa124 | 223 | 4 | 56 |
| *Pseudomonas aeruginosa* | NZ-CP008864 W6085 | 223 | 4 | 56 |
| *Pseudomonas aeruginosa* | NC-002516.2 PAO1 | 223 | 4 | 56 |
| *Salmonella enterica* | D6322 NRRL B-4212 | 979 | 7 | 140 |
| *Staphylococcus aureus* | D6322 NRRL B-41012 | 529 | 6 | 88 |

**Table 13.** Strains and Abundance-related data from 23S rRNA gene ASVs of the primary D6322 dataset.

| Species | Strain | Raw Counts | Num. Ops | CellAbund |
| --- | --- | --- | --- | --- |
| *Bacillus subtilis* | D6322 NRRL B-354 | 374 | 10 | 37 |
| *Enterococcus faecalis* | D6322 NRRL B-537 | 583 | 4 | 146 |
| *Enterococcus faecalis* | NZ-CP008816 ATCC-29212 | 376 | 4 | 94 |
| *Enterococcus faecalis* | NZ-CP060796 EF-2001 | 207 | 4 | 52 |
| *Enterococcus faecalis* | NC-017312.1 62 | 363 | 4 | 91 |
| *Enterococcus faecalis* | NC-004668.1 V583 | 363 | 4 | 91 |
| *Enterococcus faecalis* | NZ-CP098743 M9 | 207 | 4 | 52 |
| *Enterococcus faecalis* | NZ-CP110074-BE5 | 156 | 4 | 39 |
| *Escherichia coli* | D6322 NRRL B-1109 | 454 | 7 | 64 |
| *Listeria monocytogenes* | D6322 NRRL B-33116 | 501 | 6 | 84 |
| *Listeria monocytogenes* | NZ.LT985475_JF586 | 501 | 6 | 84 |
| *Listeria monocytogenes* | NZ_CP062129.FSL.J1-175 | 501 | 6 | 84 |
| *Listeria monocytogenes* | NZ_CP007462.4b.02-66 | 501 | 5 | 100 |
| *Listeria monocytogenes* | NC.003210p1.EGD-e | 501 | 6 | 84 |
| *Pseudomonas aeruginosa* | D6322 NRRL B-3509 | 192 | 4 | 48 |
| *Pseudomonas aeruginosa* | NZ_CP069335_E02 | 192 | 4 | 48 |
| *Salmonella enterica* | D6322 NRRL B-4212 | 727 | 7 | 103 |
| *Salmonella enterica* | NC.006905.1 | 110 | 7 | 16 |
| *Staphylococcus aureus* | D6322 NRRL B-41012 | 615 | 6 | 102 |

Strains, their rawcounts, number of rRNA genes, and cellular abundance-related data from 23S rRNA gene ASVs of the primary D6322 in-silico Amplicon Dataset. Note that, where there are multiple strains of a species, the data given assumes all reads associated with relevant ASVs are entirely allocated to each strain. Apportioning is addressed in the merging process.

**Table 14.** Strains and Cellular Abundance estimates following merging - primary D6322, both rRNA genes.

| Species | Strain | 16S rRNA gene CellAbund | 23S rRNA gene Cell Abund |
| --- | --- | --- | --- |
| B.subtilis | D6322 NRRL B-354 | 56 | 37 |
| E.faecalis | D6322 NRRL B-537 | 118, 0, 0 | 134, 0, 0 |
| E.faecalis | NZ-CP008816-29212 | 0, 118, 118 | 0, 94, 94 |
| E.faecalis | NZ-CP060796 EF-2001 | -, -, - | -, -, - |
| E.faecalis | NC-017312.1 62 | 0, 118, 0 | 0, 91, 0 |
| E.faecalis | NC-004668.1 V583 | 0, 0, 118 | 0, 0, 91 |
| E.faecalis | NZ-CP098743 M9 | -, -, - | -, -, - |
| E.faecalis | NZ-CP110074-BE5 | 118, 0, 0 | 52, 0, 0 |
| E.coli | D6322 NRRL B-1109 | 68 | 65 |
| L.monocytogenes | D6322 NRRL B-33116 | 133 | 84 |
| P.aeruginosa | D6322 NRRL B-3509 | 56 | 48 |
| S.enterica | D6322 NRRL B-4212 | 140 | 120 |
| S.aureus | D6322 NRRL B-41012 | 88 | 102 |

Strains and Cellular Abundance estimates from merged 16S rRNA gene and 23S rRNA gene analyses of the D6322full dataset. Note that E.faecalis has residual ambiguity of 3, and the strains listed for this species are all shared. The cellular abundances of each strain in the 3 doublets of strains for E.faecalis are listed for both 16S after merging and 23S after merging. See the illustrative examples of Methods 2.6 for detail on dealing with strain ambiguity, and Case 3 for this particular E.faecalis data.

In these tables there are entries for strains that contain the text D6322. This indicates the species-relevant D6322 strain, as downloaded from the Zymo Research website [10]. This shows that 4 species have a single entry, and in each case this corresponds to the mock microbiome strain. All 3 of the species that have multiple strains identified - *Enterococcus faecalis*, *Escherichia coli* and *Pseudomonas aeruginosa* - include the relevant D6322 strain.

It will be noted that there are only 25 entries in Table 12 despite 101 ASVs being returned by RAD denoising. It is found that there are multiple sets of ASVs such that the various ASVs in any one set have exactly the same strains that give the optimal alignment PID. There is no constraint on what the value of this optimal PID is for any ASV, but it was found to be very close to 100 (usually corresponding to a single mis-match or gap, and always less than 5 - see sub-Section 3.1). In this case there are 25 such ASV sets for the 16S rRNA gene and 20 for the 23S rRNA gene.

Table 13 gives corresponding results for the 23S rRNA gene. Three species have a single strain identified, while four have multiple strains, but again each species includes the D6322 strain amongst these unresolved strains. E.faecalis has 7 strains identified as possibly present, *Listeria monocytogenes* has 5 such strains, while *Pseudomonas aeruginosa* has 2. We refer to this multiplicity of possible strains as being initial ambiguity, with the term initial distinguishing it from what we later will refer to as residual ambiguity.

These 16S rRNA gene and 23S rRNA gene results clearly indicate that the ASVs generated are accurate to the strain level. Ambiguity usually arises because the subunit sequences of other strains of the same species are identical to those of the true species. It may also arise if the ASV does not exactly match any strain but is equi-distant from sequences of two different strains. This gives optimal alignments with a small number of mismatches or gaps, as seen in sub-Section 3.1.

The following sub-section addresses quantification, to which columns 3 - 5 in Table 12 and Table 13 relate. Sub-section 3.5 presents results from merging the 16S rRNA gene and 23S rRNA gene strain level data.

### 3.4 Cellular Abundance Estimation

RAD returns the nucleotide sequence for each ASV it determines, and also a count of reads for each ASV. Any read is associated with exactly one ASV. We are interested in the number of reads that are most closely associated with a particular strain. The ASVs of a single ASV set described above all have the same relation to strains. Hence the sum of the counts from each ASV of a single ASV set is what is given in Table 12 as the Raw Count for a strain. Where there is ambiguity in strain for a particular ASV set, all strains having the optimal alignment are listed. Also listed is the number of 16S rRNA gene or 23S rRNA gene, as appropriate, that the particular strain has. In this case all E.faecalis strains listed have the same number, as is the case for *Escherichia coli* and *Pseudomonas aeruginosa*. This is not the case for every species.

The final column of Table 12 gives the derived (relative) count for cells - it is assumed that all operons of a particular strain are equally likely to be sequenced and hence the cellular abundance is simply the total count of rRNA genes of that strain divided by the number of operons (the term operon is being used here as a generic term to describe the full ribosomal RNA operon, or any particular sub-unit, such as the 16S rRNA gene, 23S rRNA gene, V2-V3 hypervariable region etc.).

Consider now the 23S rRNA gene. For this dataset it introduces some further considerations in the analysis.

Table 13 shows there is initial ambiguity for species *Enterococcus faecalis, Listeria monocytogenes, Pseudomonas aeruginosa* and *Salmonella enterica*. The consequence is considerable variation in the cellular abundances, with different reasons for the different species. For *Enterococcus faecalis* there is a difference in the raw counts. This arises from the fact that two strains - e.g. D6322 NRRL B-537 and NZ-CP060796 EF-2001 - have different ASV sets returning these strains as optimal alignments. The 110 ASVs identified by RAD for the 23S rRNA gene reads fall into 20 ASV sets. For all of the ambiguous *Listeria monocytogenes* strains there is a single ASV set consisting of ASVs 13, 14, 16, 36, 69, 70, 99, 102, and 108. Consequently the raw count is the same for all these strains. The cellular abundance counts, however, differ by strain as the number of operons per strain differs.

For *Enterococcus faecalis*, however, different ASV sets - for instance the set of ASVs (1, 8, 61, 76) align optimally to the NZ CP008816 ATCC 29212 strain, and give a raw count of 497; the set (5, 7, 54, 64) is the only one for strains NZ CP060796 EF-2001, and NZ CP098743 M9, with a raw count of 278; the set(1, 8, 61, 76), as well as the set (5, 7, 54, 64) align optimally to strain D6322 NRRL B-537, giving a raw count of 775(=497+278). Thus we see that the raw counts for these strains reflect the ASV set with which they are associated. And because the number of operons is 4 for all these strains of *Enterococcus faecalis*, the cellular abundances estimates for strains are not needed for species cellular abundance calculation.

*Pseudomonas aeruginosa* has two strains giving ambiguity. Both have the same ASVs giving optimal alignments (8, 98) and have the same number of operons and hence the cellular abundance estimates are identical. *Salmonella enterica* is an interesting case when the 23S rRNA gene is being used for analysis. For a significant proportion of *Salmonella enterica* strains the lengths of operons within the strain fall into two groups with the lengths very similar within the group but differing by about 100 bases between the groups. For the D6322 strain, 1 operon is markedly longer than the others. Noting that strain NC 006905.1 is identified with the single ASV set of ASVs (15, 41, 63, 68, 96, 97), and that the raw count is approximately a sixth of that for D6322 NRRL B-4212, it is probable that the edit distance between some operon, or operons, of this strain and the long operon of the D6322 strain is very small.

These results, specifically for the *Listeria monocytogenes* strains, show an often-disregarded matter associated with species proportions estimation in amplicon-based microbiota analysis - cellular abundance derived from amplicon counts depends on a knowledge of which strains of some species are present, since the number of operons differs between strains. Vetrovsky and Baldrian ([11] have noted this in the context of 16S rRNA gene-based analyses. Data showing this variability for 16S rRNA gene is available through the rrnDB of Stoddard et al ([12]), but corresponding data for the 23S rRNA gene does not appear to have been developed, though it is likely to be almost identical - if not identical - to that for the 16S rRNA gene.

### 3.5 Merged 16S rRNA gene and 23S rRNA gene results

Merging of results from the 16S rRNA gene and 23S rRNA gene analyses is expected to reduce uncertainty about the number of strains, and hence perhaps improve abundance estimates. For each species the intersection of the strains was identified. Section 2.6 gives details on the principles of merging used here, rules used in implementing those principles, and some illustrative examples. The merging process involves a decision-making process that identifies one or more minimal sets of strains that meet the rules introduced in Section 2.6.1. The term residual ambiguity for a species is used for the number of such sets identified. Table 14 presents the identification and cellular abundance resulting from this process. For the primary D6322 dataset merging results in a unique strain for 6 of the species (residual ambiguity=1). However the 23S data shows *Enterococcus faecalis* remains with 5 strains which lie in one or more of the 3 doublet sets of strains (initial ambiguity=5, residual ambiguity=3). Again, this ambiguity can be traced to the very small distances between many of the E.faecalis strains at both the 16S rRNA gene and 23S rRNA gene level. Overall, as expected, residual ambiguity decreases when merging is used.

It might be expected that the sub-sampled datasets would tend to give more ambiguity. The four sub-sampled datasets detailed here provide little support for this conjecture - see Table 15. However the number of ASVs generated certainly decreases with sub-sampling.

**Table 15.** Residual ambiguity after merging for each dataset.

| Dataset | Bs | Ef | Ec | Lm | Pa | Se | Sa | Bs | Ef | Ec | Lm | Pa | Se | Sa |
| --- | --- | --- | --- | --- | --- | --- | --- | --- | --- | --- | --- | --- | --- | --- |
| D6322 | 1 | 3 | 1 | 1 | 1 | 1 | 1 | 1 | 3 | 1 | 1 | 1 | 1 | 1 |
| sub1 | 1 | 3 | 1 | 1 | 1 | 1 | 1 | 1 | 3 | 1 | 1 | 1 | 1 | 1 |
| sub2 | 1 | 10 | 1 | 1 | 1 | 1 | 1 | 1 | 0 | 1 | 1 | 1 | 1 | 1 |
| sub3 | 1 | 5 | 1 | 1 | 1 | 1 | 1 | 1 | 5 | 1 | 1 | 1 | 1 | 1 |
| sub4 | 1 | 3 | 1 | 2 | 1 | 1 | 1 | 1 | 3 | 1 | 2 | 1 | 1 | 1 |

Residual ambiguity of strains for each species in the different sets after merging 16S rRNA gene and 23S rRNA gene identification results to disambiguate. Left side for 16S rRNA gene, right side for 23 rRNA gene. Note that, for dataset sub2, no 23S ASV optimally aligned with any *E.faecalis* reference sequence, and hence merging could not be implemented and so the 16S residual ambiguity remains high. The 23S residual ambiguity for *E.faecalis* has been entered as zero.

To give some quantification of the within-species dis-ambiguation effect the following analysis is provided, with results in Table 16. For each species we determine the difference in residual ambiguity before merging and after merging for both the 16S rRNA gene and the 23S rRNA gene - see Table 16 results.

**Table 16.** Reduction in residual ambiguity from merging 16S and 23S data.

| Species | disAmbig16 | disAmbig23 |
| --- | --- | --- |
| *P.aeruginosa* | 40 | 0 |
| *S.aureus* | 4 | 0 |
| *E.coli* | 0 | 0 |
| *S.enterica* | 0 | 0 |
| *E.faecalis* | 26 | 0 |
| *L.monocytogenes* | 0 | 19 |
| *B.subtilis* | 0 | 1 |

Differences between pre-merged and post-merged 16S and 23S residual ambiguity for each species, summed across all 5 datasets.

It is noted that some disambiguation occurs for 5 of 7 species, with the effect being quite large, in most cases, for those species with high ambiguity.

Figure 3 presents barplots of strain proportions for the primary D6322 and the sub1 datasets after merging. The sub1 dataset is chosen simply as an example of the sub-sampled datasets. There remains some residual ambiguity of strains for some species. Since such residual ambiguity has been resolved by a random selection of still-ambiguous strains (Section 2.6.1), then, if such strains within a species vary in the number of operons, cellular abundances will vary. Therefore the proportions can change by a small to moderate amount from one estimation to another. For the primary D6322 case a theoretical set of proportions is shown based on the original sample being constructed from equal masses of genomic DNA. We have no guarantee that these proportions were accurately represented in the pre- or post-filtered in-silico amplicon datasets. In this context, theoretical only applies for the primary D6322 datasets. Design refers to the proportions arising when the sub-sampling factors of a particular sub-sample - see Table 10 - are applied to the estimated primary D6322 16S rRNA gene or 23S rRNA gene results. Des16, for instance, gives the proportions based on the designed sub-sampling of the primary D6322 16S rRNA gene estimated proportions. Des16 and Des23, therefore, vary according to which sub-sampled dataset is being considered. If it is accepted that the sequences from each species present are accurately attributed to species in the processing of D6322full, the proportions of each species in the sub-sampled datasets are accurately known.

**Fig. 3.**
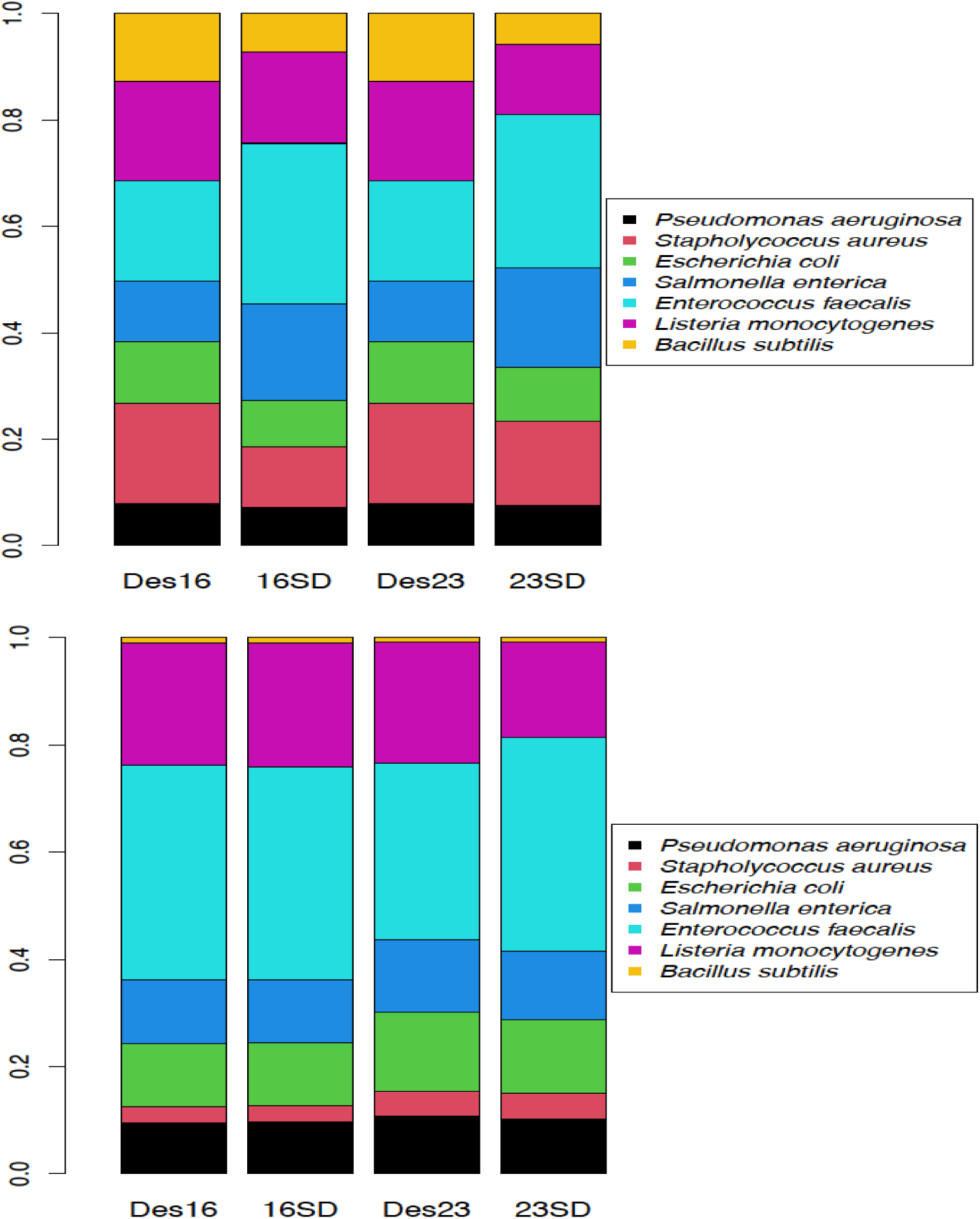
Comparison of expected (Theory, Des16, Des23) and observed (16SD, 23SD) proportions determined from 16S rRNA gene and 23S rRNA gene data after merging of 16S rRNA gene and 23S rRNA gene to reduce strain ambiguity. Results are given for the primary D6322 dataset (upper) and the sub-sampled sub01 dataset (lower). The leftmost bar gives (upper)the expected proportions based on equal masses of bacterial DNA, and (lower) the observed sub-sampled proportions. The second columns are proportions derived from the 16S rRNA gene data (after disambiguation). Columns 3 and 4 give corresponding results from the 23S rRNA gene data. The Aitchison distance between the observed proportions and the corresponding expected proportions are 1.03 (D6322, 16S), 1.09 (D6322, 23S), 0.08 (sub01, 16S) and 0.32 (sub02, 23S)

The Aitchison distances [13], [14] are distances of sets of proportions from the relevant reference proportions. - in the lower barplot of Figure 3 the value of 0.08 is the a-metric distance of 16SD from Des16. Barplots for all sub-samplings are provided in the Supplementary Material. Distances for 16S rRNA gene and 23S rRNA gene proportions were calculated for each dataset, for both pre-merged and post-merged conditions, Table 17.

**Table 17.** Distance between estimated proportions and theoretical or design proportions.

| Dataset | single16S | single23S | merged16S | merged23S |
| --- | --- | --- | --- | --- |
| D6322full | 1.04 | 1.10 | 1.04 | 1.08 |
| sub1 | 0.88 | 0.58 | 0.99 | 0.68 |
| sub2 | 0.70 | 0.21 | 0.70 | 0.21 |
| sub3 | 0.81 | 0.26 | 0.90 | 0.26 |
| sub4 | 1.48 | 1.88 | 0.92 | 0.69 |

Aitchison a-metric values for distance between estimated proportions and theoretical or design proportions. Theoretical only applies for D6322full datasets. Design refers to the proportions arising when the sub-sampling factors of a particular sub-sample are applied to the estimated D6322full 16S rRNA gene or 23S rRNA gene results.

Because the primary D6322 datasets do not give identical species proportions but form the base for sub-sampling, the pairs of 16S rRNA gene and 23S rRNA gene sub-sampled datasets are not expected to give the same proportions for the species. Therefore we cannot merge abundance proportions across 16S rRNA gene and 23S rRNA gene. With a real dataset such averaging of proportions would be valid.

## 4 Discussion

The methods and results presented here show that smart denoising of 16S rRNA gene and 23S rRNA gene, nanopore-sequenced, amplicon data of bacterial microbiomes has great potential to give near-strain, or better, taxonomic resolution of such microbiomes. We have introduced principles and methods for processing the output of a state-of-the-art denoiser, RAD, that are novel and effective in processing the current nanopore-sequenced microbial data to deal with strain ambiguity - whether that ambiguity arises from imperfect amplicon sequence variants or cross-strain identicality of amplicon sequences. We have developed a method for merging results from independent 16S rRNA gene and 23S rRNA gene analyses that achieves marked reduction in strain ambiguity within species, and suggests that cellular abundance estimates for the microbiota concerned are improved. The sub-sampling method of generation of more demanding datasets than the D6322 “even” mock microbiota has provided deeper insight into the sensitivity of the methods developed, and the power of the RAD denoiser. Overall, this work has provided a basis for much wider use of low-cost, long-read, amplicon-based nanopore sequencing to characterise bacterial microbiota at high taxonomic resolution and with improved species-level and below relative abundance estimation.

The alignment data clearly shows that under certain circumstances there is resolution between rRNA genes of a given strain. This is most obvious with the *Salmonella enterica* strain where, for the 23S rRNA genes there are 6 that are quite close and 1 which is at a Levenshtein distance greater than 100 to the others. (Supplementary Material Table 10). Multiple 23S ASVs aligned to genes indexed as 1 to 6 of the D6322 strain, while another set of 23S ASVs aligned only to a different strain. The counts of the associated reads for these two groupings, 934 and 160 respectively (Table 13) were in a close to 6:1 ratio, as would be expected. However, note that for the 16S rRNA gene of this strain, there is no strain ambiguity, and the 16S rRNA gene 7 is distant 12 from the other genes of that strain. Based on the genomes considered in this work, the distance between rRNA genes of a single strain are very often too close to be resolved by ASVs generated from the current quality of nanopore sequencing. This precludes the general use of intra-strain rRNA gene variations to assist in ambiguity reduction. The primary limitation of the work lies in not having real microbiota samples that have multiple within-species strains on which to evaluate the methods. Such a sample would need to have been characterised by alternate methods that can reliably resolve to the strain level, as well as there being an adequate ONT R10.4.1 metagenome dataset of 16S rRNA gene and 23S rRNA gene dataset generated. In the absence of such data this analysis, nevertheless, provides a set of methods, and a strong indication of their potential, for markedly improved capability from the use of nanopore technology in microbiota analyses.

## Declarations

- Funding: ZZ had an Alan W Harris Scholarship from WEHI while undertaking his Master of Science research project.
- Conflict of interest. Not applicable.
- Ethics approval. Not applicable.
- Consent to participate. Not applicable.
- Consent for publication. Not applicable.
- Availability of data and materials

- The metagenomic dataset supporting the conclusions of this article is available at https://www.ebi.ac.uk/ena/browser/view/PRJEB48692 and was accessed with the following Unix call wget ftp://ftp.sra.ebi.ac.uk/vol1/runERR728/ERR7287988/zymo hmw r104.fastq.gz, giving a 48 Gb file.
- Sequences of 16S and 23S rRNA genes extracted from that 48Gb file gave files amplicon 16S D6322 trimmed 05022024.fastq and amplicon 23S D6322 trimmed 05022024.fastq. These are used as input to RAD - approx. 67MB and 104 MB fastq files that need to be placed in a repository for access.
- Additional Files are on Figshare at https://melbourne.figshare.com/account/home/projects/195149. These consist of a Supplementary Material file, 9 tarred files of input adn output data, together with an explanatory text file.
- ASVs generated for the primary D6322 16S rRNA gene and 23S rRNA gene - provided as fasta files in Additional Files
- Sub-sampled datasets, both 16S rRNA gene and 23S rRNA gene as input to RAD
- provided as fastq files as Additional Files.
- ASVs generated for each sub-sampled dataset - provided as fasta files in Additional Files
- Fasta files from which 16S rRNA gene and 23S rRNA gene reference DBs were generated - provided as fasta files in Additional Files.
- Code availability - Github repository https://github.com/cjwoodruff50/Nanopore WEHI cjw.
- Authors’ contributions

- CJW conceived the study, developed methods, coded and interpreted the analyses, provided supervision, and wrote the manuscript.
- ZZ analysed bacterial 23S rRNA gene sequences to identify hypervariable regions and defined and in-silico validated these.
- TPS provided critical assessment and statistical insight at all stages of method and analysis development, and assisted in supervision, and manuscript preparation.

## Acknowledgments

Valuable support in using the RAD denoising code and the visualisation code, seqUMAP, was provided by Benjamin Murrell, Department of Microbiology, Tumor and Cell Biology, Karolinska Institutet, Sweden.

