## SupplementaryMaterial for "Full 16S and 23S rRNA gene-based,strain-level resolution of the microbiota of a mock bacterial microbiome using ONT nanopore sequencing"

### Microbiome ASV Supplementary Material 08022024

February 2024

*[1]Christopher J. Woodruff
[1]Zhengming Zhang
*[1]Terence P. Speed
*[1]Bioinformatics Division, WEHI, 1G Royal Parade, Parkville, 3052, Victoria, Australia

#### 1 Strain Level Matching Detail

A set of randomly selected strains was created for each of the 7 species of the D6322 mock microbiota. The intent was to have a challenging strain-level identification task for most species. It is considered that, if the maximum Levenshtein distance between all pairs of operons from a pair of strains is small, then discrimination of these strains will be difficult. Therefore, for each species the Levenshtein distances between all pairs of operons of each pair of strains were calculated. These pairwise strain inter-operon distances can be represented as matrices which provide a detailed representation of how similar two strains are. For each species a set of such strain-pair matrices arises which can be represented as a symmetric block matrix. Below we present tabular data for each species giving the intra-strain distances between 16S rRNA genes for just the D6322 strain, and likewise for the 23S rRNA genes. Each such table would form a single block on the diagonal in one of the 14 symmetric block - 7 species, 2 rRNA genes - matrices described.

*Bacillus subtilis* , D6322 strain has 10 operons. Table 1 and 2 show the distances between *Bacillus subtilis* 16S rRNA genes of the D6322 strain, while Table 2 gives corresponding data for the 23S rRNA gene. All are small, and it can be seen that operons 2 and 6 are identical, as are 3 and 7, and 4 and 5.

Tables 3 to 12 give corresponding data for the D6322 strain of the other species.

No tables are included for *Pseudomonas aeruginosa* as all 16S rRNA genes and all 23 rRNA genes are identical for the D6322 strain.

Table 1: *Bacillus subtilis* D6322 strain pairwise 16S rRNA gene separations.

| Operon | Op 1 | Op 2 | Op 3 | Op 4 | Op 5 | Op 6 | Op 7 | Op 18 | Op 9 | Op 10 |
|---|---|---|---|---|---|---|---|---|---|---|
| Op 1 | 0 | 29 | 2 | 1 | 1 | 29 | 29 | 31 | 30 | 2 |
| Op 2 | 29 | 36 | 35 | 35 | 3 | 3 | 3 | 2 | 35 | 2 |
| Op 3 | 2 | 32 | 0 | 1 | 1 | 36 | 36 | 36 | 35 | 2 |
| Op 4 | 1 | 31 | 1 | 0 | 0 | 35 | 35 | 35 | 34 | 1 |
| Op 5 | 1 | 31 | 1 | 0 | 0 | 35 | 35 | 35 | 34 | 1 |
| Op 6 | 29 | 3 | 36 | 35 | 35 | 0 | 0 | 2 | 3 | 35 |
| Op 7 | 29 | 3 | 36 | 35 | 35 | 0 | 0 | 2 | 3 | 35 |
| Op 8 | 31 | 3 | 36 | 35 | 35 | 2 | 2 | 0 | 3 | 35 |
| Op 9 | 30 | 2 | 35 | 34 | 34 | 3 | 3 | 3 | 0 | 34 |
| Op 10 | 2 | 31 | 2 | 1 | 1 | 35 | 35 | 35 | 34 | 0 |

Table 2: *Bacillus subtilis* D6322 strain pairwise 23S rRNA gene separations.

| Operon | Op 1 | Op 2 | Op 3 | Op 4 | Op 5 | Op 6 | Op 7 | Op 18 | Op 9 | Op 10 |
|---|---|---|---|---|---|---|---|---|---|---|
| Op 1 | 0 | 2 | 1 | 3 | 3 | 2 | 1 | 3 | 3 | 4 |
| Op 2 | 2 | 0 | 1 | 1 | 1 | 0 | 1 | 1 | 1 | 2 |
| Op 3 | 1 | 1 | 0 | 2 | 2 | 1 | 0 | 2 | 2 | 3 |
| Op 4 | 3 | 1 | 2 | 0 | 0 | 1 | 2 | 2 | 2 | 1 |
| Op 5 | 3 | 1 | 2 | 0 | 0 | 1 | 2 | 2 | 2 | 1 |
| Op 6 | 2 | 0 | 1 | 1 | 1 | 0 | 1 | 1 | 1 | 2 |
| Op 7 | 1 | 1 | 0 | 2 | 2 | 1 | 0 | 2 | 2 | 3 |
| Op 8 | 3 | 1 | 2 | 2 | 2 | 1 | 2 | 0 | 2 | 3 |
| Op 9 | 3 | 1 | 2 | 2 | 2 | 1 | 2 | 2 | 0 | 3 |
| Op 10 | 4 | 2 | 3 | 1 | 1 | 2 | 3 | 3 | 3 | 0 |

Table 3: *Enterococcus faecalis* D6322 strain pairwise 16S rRNA gene separations.

| Operon | Op 1 | Op 2 | Op 3 | Op 4 |
|---|---|---|---|---|
| Op 1 | 0 | 0 | 0 | 0 |
| Op 2 | 0 | 0 | 0 | 0 |
| Op 3 | 0 | 0 | 0 | 0 |
| Op 4 | 0 | 0 | 0 | 0 |

Table 4: *Enterococcus faecalis* D6322 strain pairwise 23S rRNA gene separations.

| Operon | Op 1 | Op 2 | Op 3 | Op 4 |
|---|---|---|---|---|
| Op 1 | 0 | 1 | 2 | 4 |
| Op 2 | 1 | 0 | 1 | 3 |
| Op 3 | 2 | 1 | 0 | 4 |
| Op 4 | 4 | 3 | 4 | 0 |

Table 5: *Escherichia coli* D6322 strain pairwise 16S rRNA gene separations.

| Operon | Op 1 | Op 2 | Op 3 | Op 4 | Op 5 | Op 6 | Op 7 |
|---|---|---|---|---|---|---|---|
| Op 1 | 0 | 8 | 8 | 0 | 8 | 8 | 3 |
| Op 2 | 8 | 0 | 0 | 8 | 0 | 0 | 2 |
| Op 3 | 8 | 0 | 0 | 8 | 0 | 0 | 2 |
| Op 4 | 0 | 8 | 8 | 0 | 8 | 8 | 3 |
| Op 5 | 8 | 0 | 0 | 8 | 0 | 0 | 2 |
| Op 6 | 8 | 0 | 0 | 8 | 0 | 0 | 2 |
| Op 7 | 3 | 2 | 2 | 3 | 2 | 2 | 0 |

Table 6: *Escherichia coli* D6322 strain pairwise 23S rRNA gene separations.

| Operon | Op 1 | Op 2 | Op 3 | Op 4 | Op 5 | Op 6 | Op 7 |
|---|---|---|---|---|---|---|---|
| Op 1 | 0 | 0 | 0 | 0 | 0 | 0 | 0 |
| Op 2 | 0 | 0 | 0 | 0 | 0 | 0 | 0 |
| Op 3 | 0 | 0 | 0 | 0 | 0 | 0 | 0 |
| Op 4 | 0 | 0 | 0 | 0 | 0 | 0 | 0 |
| Op 5 | 0 | 0 | 0 | 0 | 0 | 0 | 0 |
| Op 6 | 0 | 0 | 0 | 0 | 0 | 0 | 0 |
| Op 7 | 0 | 0 | 0 | 0 | 0 | 0 | 0 |

Table 7: *Listeria monocytogenes* D6322 strain pairwise 16S rRNA gene separations.

| Operon | Op 1 | Op 2 | Op 3 | Op 4 | Op 5 | Op 6 |
|---|---|---|---|---|---|---|
| Op 1 | 0 | 1 | 2 | 1 | 1 | 1 |
| Op 2 | 1 | 0 | 1 | 0 | 0 | 0 |
| Op 3 | 2 | 1 | 0 | 1 | 1 | 1 |
| Op 4 | 1 | 0 | 1 | 0 | 0 | 0 |
| Op 5 | 1 | 0 | 1 | 0 | 0 | 0 |
| Op 6 | 1 | 0 | 1 | 0 | 0 | 0 |

Table 8: *Listeria monocytogenes* D6322 strain pairwise 23S rRNA gene separations.

| Operon | Op 1 | Op 2 | Op 3 | Op 4 | Op 5 | Op 6 |
|---|---|---|---|---|---|---|
| Op 1 | 0 | 0 | 0 | 0 | 1 | 3 |
| Op 2 | 0 | 0 | 0 | 0 | 1 | 3 |
| Op 3 | 0 | 0 | 0 | 0 | 1 | 3 |
| Op 4 | 0 | 0 | 0 | 0 | 1 | 3 |
| Op 5 | 1 | 1 | 1 | 1 | 0 | 4 |
| Op 6 | 3 | 3 | 3 | 3 | 4 | 0 |

Table 9: *Salmonella enterica* D6322 strain pairwise 16S rRNA gene separations.

| Operon | Op 1 | Op 2 | Op 3 | Op 4 | Op 5 | Op 6 | Op 7 |
|---|---|---|---|---|---|---|---|
| Op 1 | 0 | 0 | 0 | 0 | 0 | 0 | 12 |
| Op 2 | 0 | 0 | 0 | 0 | 0 | 0 | 12 |
| Op 3 | 0 | 0 | 0 | 0 | 0 | 0 | 12 |
| Op 4 | 0 | 0 | 0 | 0 | 0 | 0 | 12 |
| Op 5 | 0 | 0 | 0 | 0 | 0 | 0 | 12 |
| Op 6 | 0 | 0 | 0 | 0 | 0 | 0 | 12 |
| Op 7 | 30 | 30 | 12 | 12 | 12 | 12 | 0 |

Table 10: *Salmonella enterica* strain pairwise 23S rRNA gene separations.

| Operon | Op 1 | Op 2 | Op 3 | Op 4 | Op 5 | Op 6 | Op 7 |
|---|---|---|---|---|---|---|---|
| Op 1 | 0 | 5 | 3 | 6 | 2 | 3 | 105 |
| Op 2 | 5 | 0 | 4 | 7 | 5 | 4 | 102 |
| Op 3 | 3 | 4 | 0 | 5 | 3 | 2 | 106 |
| Op 4 | 6 | 7 | 5 | 0 | 6 | 5 | 109 |
| Op 5 | 2 | 5 | 3 | 6 | 0 | 1 | 105 |
| Op 6 | 3 | 4 | 2 | 5 | 1 | 0 | 106 |
| Op 7 | 105 | 102 | 106 | 109 | 105 | 106 | 0 |

Table 11: *Staphylococcus aureus.* strain pairwise 16S rRNA gene separations.

| Operon | Op 1 | Op 2 | Op 3 | Op 4 | Op 5 | Op 6 |
|---|---|---|---|---|---|---|
| Op 1 | 0 | 3 | 1 | 0 | 0 | 0 |
| Op 2 | 3 | 0 | 4 | 3 | 3 | 3 |
| Op 3 | 1 | 4 | 0 | 1 | 1 | 1 |
| Op 4 | 0 | 3 | 1 | 0 | 0 | 0 |
| Op 5 | 0 | 3 | 1 | 0 | 0 | 0 |
| Op 6 | 0 | 3 | 1 | 0 | 0 | 0 |

Table 12: *Staphylococcus aureus.* D6322 strain pairwise 23S rRNA gene separations.

| Operon | Op 1 | Op 2 | Op 3 | Op 4 | Op 5 | Op 6 |
|---|---|---|---|---|---|---|
| Op 1 | 0 | 1 | 1 | 20 | 0 | 1 |
| Op 2 | 1 | 0 | 0 | 21 | 1 | 2 |
| Op 3 | 1 | 0 | 0 | 21 | 1 | 2 |
| Op 4 | 20 | 21 | 21 | 0 | 20 | 21 |
| Op 5 | 0 | 1 | 1 | 20 | 0 | 1 |
| Op 6 | 1 | 0 | 0 | 21 | 1 | 2 |

#### 2 Visualisation of ASVs Based on distance between them

Based on RAD output and our variant of the Murrell Lab ([3]) seqUMAP, representations of the clustering of ASVs and the reads associated with those ASVs are provided in Figures 1 and 2. The approximate edit distance of Kumar et al. [1], given by equation (1), is used.

$$d(,) = \frac{\sum_{j=1}^{4^k}(a_j - b_j)^2}{k(||+||)} \tag{1}$$

where $k = 6$ for 6-mers and $||, ||$ are the lengths of the sequences ,. Equation 1 was used to generate inter-ASV and inter-read distance data that were then processed by UMAP to give these 2D representations of the relative positions of ASVs and reads for both 16S rRNA gene datasets and 23S rRNA gene datasets in this space.

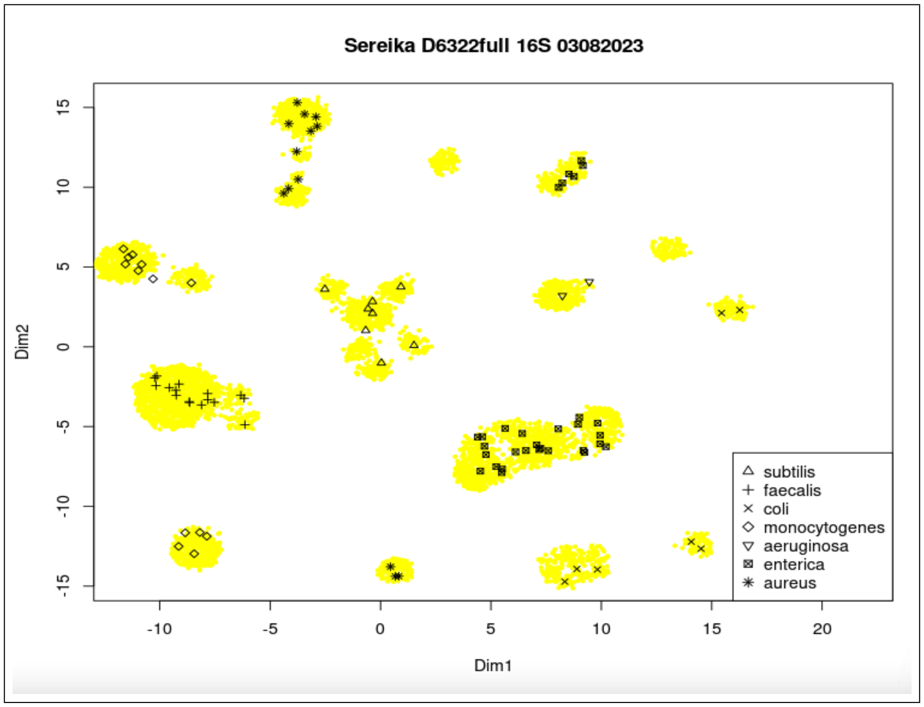


Figure 1: UMAP-derived 2D representation of 16S rRNA gene ASVs (black symbols) and reads (yellow dots) with identification of the species of the D6322 strains to which these ASVs are classified The two clusters of reads without any ASVs co-located are ASVs classified to another strain of E.coli.

Figure 1 shows ASVs associated with a particular species lying within clouds of actual 16S rRNA gene sequences, and that only a single species occurs in any of these clouds.

Figure 2 is the corresponding representation for the D6322full 23S rRNA gene reads and the ASVs associated with the various species. Here the results are not as clean, with only E.faecalis and P.aeruginosa giving read clouds with only the single species ASVs associated with them.

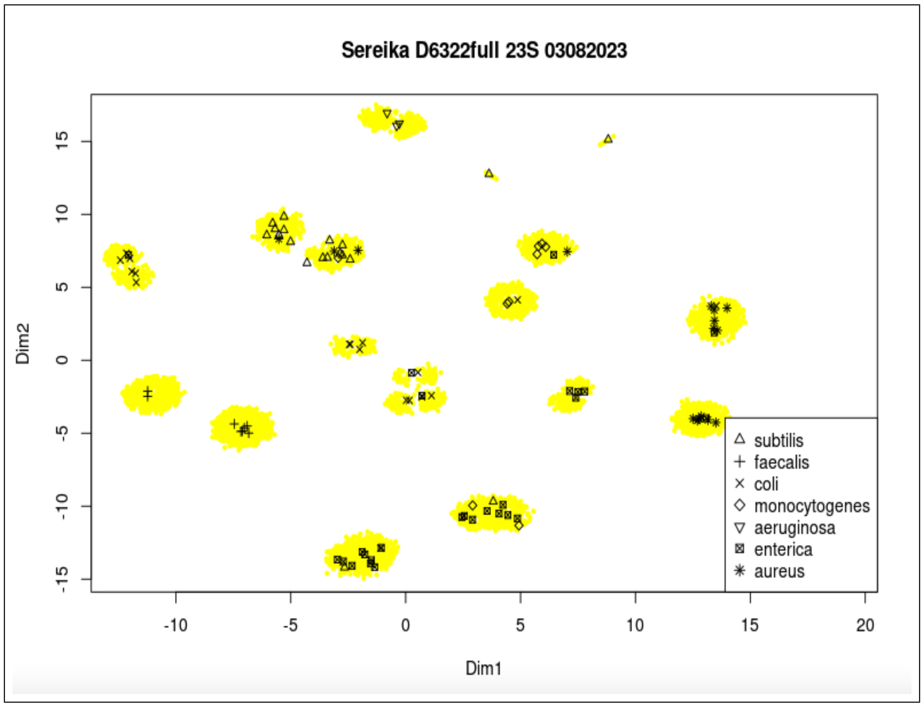


Figure 2: UMAP-derived 2D representation of 23S rRNA gene ASVs (black symbols) and reads (yellow dots) with identification of the species of the D6322 strains to which these ASVs are classified.

### 3 Quality of ASVs

Table 13 quantifies the histogram data of Figure ??. From this table it is clear that as stated in the main text, fewer than 2% of reads are associated with ASVs having more than 3 gaps plus mismatches in their optimal alignments, and more than 80% of reads are associated with ASVs differing by no more than 1 gap or 1 mismatch. One can also see the rather surprising result that the 23S rRNA gene error counts tend to be lower than the 16S rRNA gene despite the 23S rRNA gene sequences being approximately 50% longer than than the 16S rRNA gene sequences.

Table 13: Proportion of reads for various ASV alignment error rates and types

| Count | 16S rRNA gene Mismatch | 16S rRNA gene Gap | 16S rRNA gene (Gap+Mismatch) |
|---|---|---|---|
| 0 | 0.700 | 0.816 | 0.572 |
| 1 | 0.215 | 0.159 | 0.296 |
| 2 | 0.052 | 0.019 | 0.074 |
| 3 | 0.027 | 0.006 | 0.040 |
| 4 | 0.006 | 0.000 | 0.018 |

| Count | 23S rRNA gene Mismatch | 23S rRNA gene Gap | 23S rRNA gene (Gap+Mismatch) |
|---|---|---|---|
| 0 | 0.736 | 0.806 | 0.615 |
| 1 | 0.179 | 0.143 | 0.220 |
| 2 | 0.068 | 0.046 | 0.116 |
| 3 | 0.027 | 0.005 | 0.037 |
| 4 | 0.002 | 0.000 | 0.008 |

Further insight into the relation between reference sequences and ASV sequences is provided in the following tables. These give the Levenshtein distance between the 16S rRNA gene sequences of the true strain of a selected species and the ASVs that have given optimal alignments to that species., and likewise for the 23S rRNA gene sequences. The minimal distance of an ASV to an operon is also tabulated for each ASV. Tables are given for *Bacillus subtilis*, *Escherichia coli* , and *Staphylococcus aureus* - Tables 14 to 19. Similar results are obtained for the other species.
Lengths of operons: 1574 1573 1573 1573 1573 1573 1573 1573 1573 1573
Length range of ASVs: 1513 - 1515

For *Bacillus subtilis* , for both 16S and 23S rRNA genes, all but 1 of the ASVs are within a distance of 3 from an operon, with operon 52 being at a distance of 5. All but 1 of the 10 16S rRNA genes have an ASV that aligns exactly with them, and all but 2 of the 23S rRNA genes have an ASV that aligns exactly with them. Note that ASVs are reliably shorter than the relevant operons. For both sets of rRNA genes the majority of ASVs give a perfect alignment.
Lengths of operons: 2586 2583 2584 2584 2584 2584 2584 2568 2584 2584 .
Lengths range of ASVs: 2505 - 2510

Table 14: Data on how similar ASVs are to 16SrRNA genes of the true strain - *Bacillus subtilis*.

| ASV | Op1 | Op2 | Op3 | Op4 | Op5 | Op6 | Op7 | Op8 | Op9 | Op10 | BestMatch |
|---|---|---|---|---|---|---|---|---|---|---|---|
| ASV_6 | 1 | 2 | 3 | 2 | 2 | 0 | 0 | 2 | 3 | 2 | 0 |
| ASV_10 | 2 | 1 | 2 | 1 | 1 | 3 | 3 | 3 | 0 | 1 | 0 |
| ASV_14 | 2 | 1 | 0 | 1 | 1 | 3 | 3 | 3 | 2 | 1 | 0 |
| ASV_16 | 3 | 2 | 3 | 2 | 2 | 2 | 2 | 0 | 3 | 2 | 0 |
| ASV_3 | 1 | 0 | 1 | 0 | 0 | 2 | 2 | 2 | 1 | 0 | 0 |
| ASV_36 | 2 | 1 | 2 | 1 | 1 | 3 | 3 | 3 | 2 | 1 | 1 |
| ASV_46 | 2 | 1 | 2 | 1 | 1 | 3 | 3 | 3 | 2 | 1 | 1 |
| ASV_58 | 4 | 3 | 4 | 3 | 3 | 5 | 5 | 5 | 4 | 3 | 3 |

Table 15: Data on how similar ASVs are to 23S rRNA genes of the true strain - *Bacillus subtilis*.

| ASV | Op1 | Op2 | Op3 | Op4 | Op5 | Op6 | Op7 | Op8 | Op9 | Op10 | BestMatch |
|---|---|---|---|---|---|---|---|---|---|---|---|
| ASV_16 | 3 | 3 | 2 | 4 | 4 | 3 | 2 | 4 | 4 | 5 | 2 |
| ASV_24 | 1 | 1 | 0 | 2 | 2 | 1 | 0 | 2 | 2 | 3 | 0 |
| ASV_18 | 4 | 2 | 3 | 1 | 1 | 2 | 3 | 3 | 3 | 0 | 0 |
| ASV_74 | 4 | 2 | 3 | 1 | 1 | 2 | 3 | 3 | 3 | 0 | 0 |
| ASV_20 | 3 | 1 | 2 | 0 | 0 | 1 | 2 | 2 | 2 | 1 | 0 |
| ASV_31 | 3 | 1 | 2 | 0 | 0 | 1 | 2 | 2 | 2 | 1 | 0 |
| ASV_79 | 6 | 4 | 5 | 3 | 3 | 4 | 5 | 5 | 5 | 4 | 3 |
| ASV_86 | 4 | 2 | 3 | 1 | 1 | 2 | 3 | 3 | 3 | 2 | 1 |
| ASV_92 | 4 | 2 | 3 | 1 | 1 | 2 | 3 | 3 | 3 | 2 | 1 |
| ASV_25 | 3 | 1 | 2 | 2 | 2 | 1 | 2 | 2 | 0 | 3 | 0 |
| ASV_84 | 3 | 1 | 2 | 2 | 2 | 1 | 2 | 2 | 0 | 3 | 0 |
| ASV_88 | 4 | 2 | 3 | 3 | 3 | 2 | 3 | 3 | 1 | 4 | 1 |
| ASV_78 | 3 | 1 | 2 | 2 | 2 | 1 | 2 | 0 | 2 | 3 | 0 |
| ASV_52 | 5 | 8 | 9 | 9 | 9 | 8 | 9 | 9 | 9 | 10 | 5 |
| ASV_55 | 3 | 1 | 2 | 2 | 2 | 1 | 2 | 2 | 2 | 3 | 1 |
| ASV_85 | 3 | 1 | 2 | 2 | 2 | 1 | 2 | 2 | 2 | 3 | 1 |
| ASV_87 | 2 | 0 | 1 | 1 | 1 | 0 | 1 | 1 | 1 | 2 | 0 |
| ASV_98 | 2 | 0 | 1 | 1 | 1 | 0 | 1 | 1 | 1 | 2 | 0 |
| ASV_107 | 5 | 3 | 4 | 4 | 4 | 3 | 4 | 4 | 4 | 5 | 3 |

Table 16: Data on how similar ASVs are to 16S rRNA genes of the true strain - *Escherichia coli.*

| ASV | Op1 | Op2 | Op3 | Op4 | Op5 | Op6 | Op7 | BestMatch |
|---|---|---|---|---|---|---|---|---|
| ASV_31 | 13 | 8 | 9 | 10 | 0 | 10 | 13 | 0 |
| ASV_44 | 14 | 9 | 10 | 11 | 1 | 11 | 14 | 1 |
| ASV_19 | 19 | 0 | 15 | 16 | 8 | 16 | 19 | 0 |
| ASV_21 | 20 | 1 | 16 | 17 | 9 | 17 | 20 | 1 |
| ASV_35 | 12 | 16 | 1 | 2 | 10 | 2 | 5 | 1 |
| ASV_48 | 11 | 15 | 0 | 1 | 9 | 1 | 4 | 0 |
| ASV_81 | 12 | 16 | 1 | 2 | 10 | 2 | 5 | 1 |
| ASV_22 | 5 | 14 | 6 | 7 | 8 | 7 | 4 | 4 |
| ASV_68 | 6 | 15 | 7 | 8 | 9 | 8 | 5 | 5 |
| ASV_17 | 12 | 16 | 1 | 0 | 10 | 0 | 5 | 0 |
| ASV_49 | 13 | 17 | 2 | 1 | 11 | 1 | 6 | 1 |
| ASV_64 | 13 | 17 | 2 | 1 | 11 | 1 | 6 | 1 |
| ASV_82 | 14 | 18 | 3 | 2 | 12 | 2 | 7 | 2 |
| ASV_86 | 14 | 18 | 3 | 2 | 12 | 2 | 7 | 2 |
| ASV_97 | 15 | 19 | 4 | 3 | 13 | 3 | 8 | 3 |
| ASV_45 | 10 | 20 | 5 | 6 | 14 | 6 | 1 | 1 |
| ASV_78 | 9 | 19 | 4 | 5 | 13 | 5 | 0 | 0 |
| ASV_87 | 10 | 20 | 5 | 6 | 14 | 6 | 1 | 1 |

Lengths of operons: 1555 1564 1558 1564 1564 1564 1560
Length range of ASVs: 1505 -1506

For *Escherichia coli* the quality of ASV to rRNA gene matching is not as good as for *Bacillus subtilis.* Every 23S ASV has a match to a 23S rRNA gene that is no more than a distance of 3 from it, while all but 2 of 18 16S ASVs have such a match to 16S rRNA genes. Also, all 7 of the 23 rRNA genes are within a distance of 3 from an ASV, with 2 having exact matches, while for the 16S ASVs 6 of the 16S rRNA genes have an exact match to one ASV, and no ASV is more than a distance of 5 from an rRNA gene. It is worth noting that Sereika et al. found the quality of their D6322 metagenome to have lower quality of *Escherichia coli* reads than any of the other species, and our data is consistent with that observation.
Lengths of operons: 2553 2555 2557 2556 2558 2556 2557
Lengths range of ASVs: 2477 - 2482

Table 17: Data on how similar ASVs are to 23S rRNA genes of the true strain - *Escherichia coli*.

| ASV | Op1 | Op2 | Op3 | Op4 | Op5 | Op6 | Op7 | BestMatch |
|---|---|---|---|---|---|---|---|---|
| ASV_19 | 9 | 10 | 16 | 17 | 0 | 16 | 6 | 0 |
| ASV_101 | 9 | 10 | 16 | 17 | 0 | 16 | 6 | 0 |
| ASV_23 | 3 | 16 | 14 | 15 | 6 | 14 | 0 | 0 |
| ASV_28 | 3 | 16 | 14 | 15 | 6 | 14 | 0 | 0 |
| ASV_34 | 4 | 17 | 15 | 16 | 7 | 15 | 1 | 1 |
| ASV_73 | 4 | 17 | 15 | 16 | 7 | 15 | 1 | 1 |
| ASV_90 | 5 | 18 | 16 | 17 | 8 | 16 | 2 | 2 |
| ASV_35 | 17 | 28 | 2 | 3 | 16 | 2 | 14 | 2 |
| ASV_39 | 16 | 27 | 1 | 2 | 15 | 1 | 13 | 2 |
| ASV_51 | 17 | 28 | 2 | 3 | 16 | 2 | 14 | 2 |
| ASV_57 | 17 | 28 | 2 | 3 | 16 | 2 | 14 | 2 |
| ASV_65 | 16 | 27 | 1 | 2 | 15 | 1 | 13 | 2 |
| ASV_81 | 17 | 28 | 2 | 3 | 16 | 2 | 14 | 2 |
| ASV_102 | 18 | 29 | 3 | 4 | 17 | 3 | 15 | 3 |
| ASV_37 | 17 | 2 | 26 | 27 | 8 | 26 | 14 | 2 |
| ASV_59 | 18 | 3 | 27 | 28 | 9 | 27 | 15 | 3 |
| ASV_87 | 18 | 3 | 27 | 28 | 9 | 27 | 15 | 3 |
| ASV_61 | 17 | 2 | 26 | 27 | 8 | 26 | 14 | 2 |

Table 18: Data on how similar ASVs are to 16S rRNA genes of the true strain - *Staphylococcus aureus*.

| ASV | Op1 | Op2 | Op3 | Op4 | Op5 | Op6 | BestMatch |
|---|---|---|---|---|---|---|---|
| ASV_12 | 3 | 0 | 4 | 3 | 3 | 3 | 0 |
| ASV_74 | 4 | 1 | 5 | 4 | 4 | 4 | 1 |
| ASV_77 | 5 | 2 | 6 | 5 | 5 | 5 | 2 |
| ASV_5 | 0 | 3 | 1 | 0 | 0 | 0 | 0 |
| ASV_27 | 1 | 4 | 2 | 1 | 1 | 1 | 1 |
| ASV_32 | 3 | 6 | 4 | 3 | 3 | 3 | 3 |
| ASV_83 | 2 | 5 | 3 | 2 | 2 | 2 | 2 |
| ASV_88 | 2 | 5 | 3 | 2 | 2 | 2 | 2 |
| ASV_95 | 2 | 5 | 3 | 2 | 2 | 2 | 2 |
| ASV_11 | 2 | 5 | 1 | 2 | 2 | 2 | 1 |
| ASV_54 | 3 | 6 | 2 | 3 | 3 | 3 | 2 |
| ASV_84 | 4 | 7 | 3 | 4 | 4 | 4 | 4 |
| ASV_69 | 4 | 7 | 5 | 8 | 8 | 8 | 4 |

Lengths of operons: 1574 1574 1574 1575 1575 1575
Length range of ASVs: 1513 - 1517

For *Staphylococcus aureus* 4 of the 6 16S rRNA genes are identical and these have an ASV that aligns perfectly to them. Of the remaining 2 rRNA genes, one has a perfect match to an ASV and the other is distance 1 from an ASV. No 16S ASV is more distant than 4 from an operon. All but 1 of the 23S ASVs are within a distance of 3 from a 23S rRNA gene, while 5 of the 6 genes have zero distance from at least one ASV. However 1 of the 23S rRNA genes is no closer than 19 to any ASV. Reads from such an gene might be expected to "leak" reads to some ASV from another species, depending on the inter-species distances between strains.
Lengths of operons: 2582 2582 2582 2585 2583 2584
Lengths range of ASVs: 2504 - 2510

Table 19: Data on how similar ASVs are to 23S rRNA genes of the true strain - *Staphylococcus aureus.*

| ASV | Op1 | Op2 | Op3 | Op4 | Op5 | Op6 | BestMatch |
|---|---|---|---|---|---|---|---|
| ASV_6 | 1 | 0 | 0 | 21 | 1 | 2 | 0 |
| ASV_27 | 1 | 0 | 0 | 21 | 1 | 2 | 0 |
| ASV_32 | 3 | 2 | 2 | 23 | 3 | 4 | 2 |
| ASV_75 | 7 | 6 | 6 | 27 | 7 | 8 | 6 |
| ASV_77 | 2 | 1 | 1 | 22 | 2 | 3 | 1 |
| ASV_80 | 4 | 3 | 3 | 29 | 4 | 5 | 3 |
| ASV_89 | 4 | 3 | 3 | 24 | 4 | 5 | 3 |
| ASV_38 | 1 | 2 | 2 | 21 | 1 | 0 | 0 |
| ASV_109 | 2 | 3 | 3 | 22 | 2 | 1 | 1 |
| ASV_2 | 0 | 1 | 1 | 20 | 0 | 1 | 0 |
| ASV_22 | 1 | 2 | 2 | 21 | 1 | 2 | 1 |
| ASV_29 | 0 | 1 | 1 | 20 | 0 | 1 | 0 |
| ASV_56 | 3 | 4 | 4 | 19 | 3 | 4 | 3 |
| ASV_58 | 1 | 2 | 2 | 21 | 1 | 2 | 1 |
| ASV_67 | 1 | 2 | 2 | 21 | 1 | 2 | 1 |
| ASV_93 | 2 | 3 | 3 | 22 | 2 | 3 | 2 |
| ASV_108 | 2 | 3 | 3 | 22 | 2 | 3 | 2 |

#### 4 Species-level Abundance

Proportions of species for 16S rRNA gene and 23S rRNA gene for all four sub-sampled datasets are presented in Figures 3 and 4.

Note that, except for *Enterococcus faecalis* 23S for sub2, all species are present in each barplot illustrating the proportions ranging over 3 orders of magnitude.

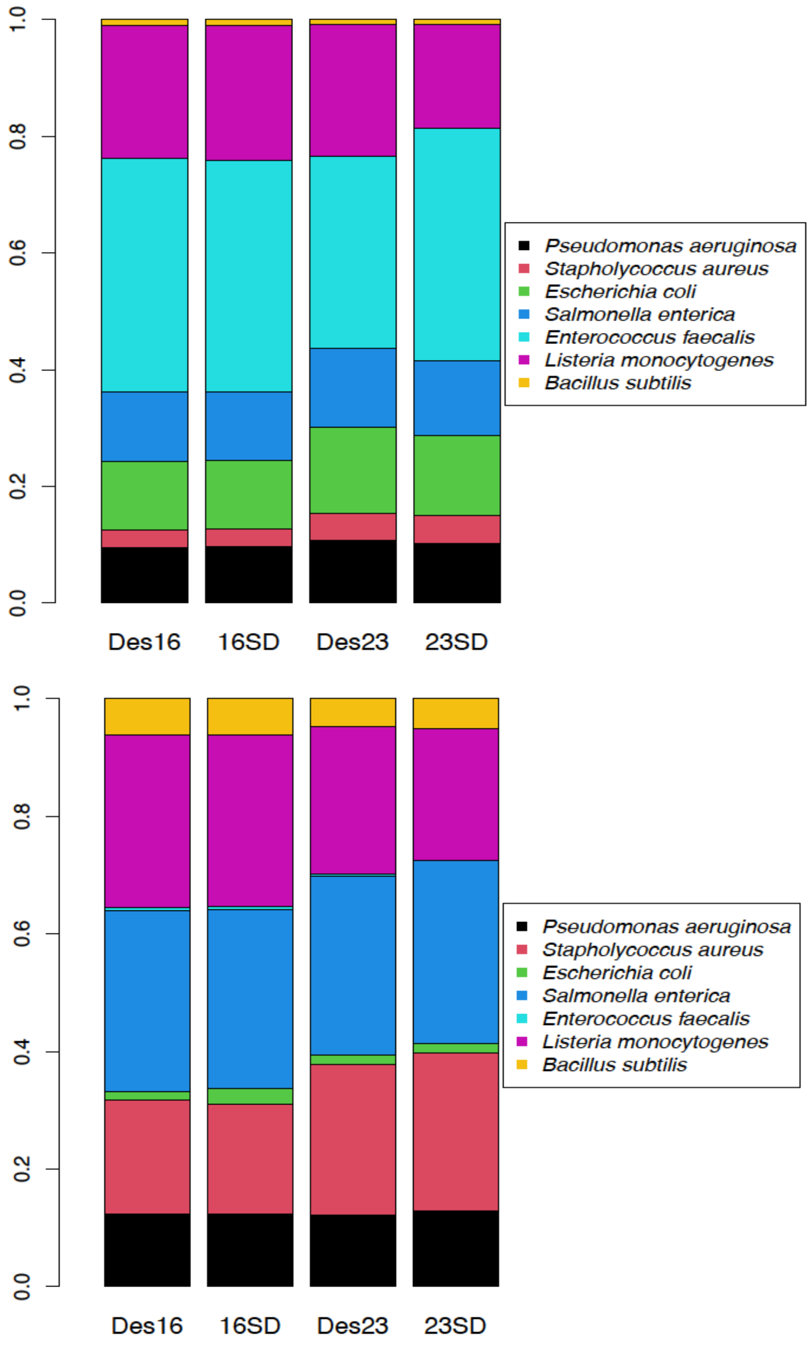


Figure 3: Comparison of expected and observed proportions for the sub1 dataset (upper) and sub2 dataset (lower). The first and third columns are the expected proportions, while the second and fourth are observed consequent on the merging of separate 16S rRNA gene and 23S rRNA gene results for sub-sampled datasets of each. The Aitchison distance between the expected proportions (Des16, Des23) and the corresponding observed proportions (16SD, 23SD) are 0.08 (sub1 16S), 0.32 (sub1 23S), 0.57 (sub2, 16S) and 0.67 (sub2 23S)

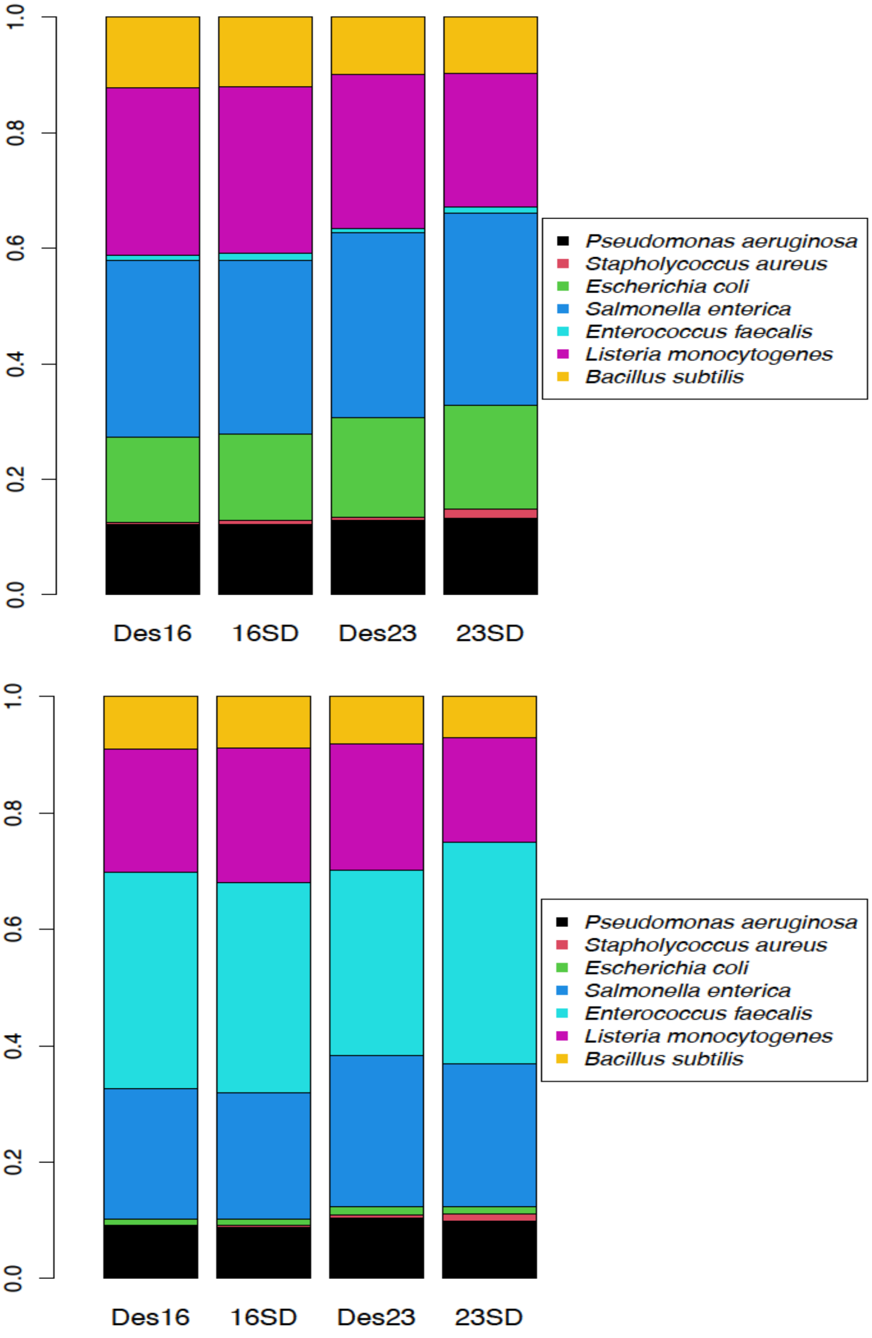


Figure 4: Comparison of expected and observed proportions for the sub3 dataset (upper) and sub4 dataset (lower). The Aitchison distance between the expected proportions (Des16, Des23) and the corresponding observed proportions (16SD, 23SD) are 0.49 (sub31 16S), 1.05 (sub3 23S), 0.14 (sub4, 16S) and 1.05 (sub4 23S)

##### 4.1 Apportioning ASV set counts across shared strains

From the rules adopted for merging 16S rRNA gene and 23S rRNA gene data within a species there is a requirement to apportion counts from an ASV set that is common to more than one of the shared strains such that the cellular abundances of each strain are equal. This was illustrated in Case 2 of subsection **??** with a specific numeric example. The following gives a general formulation of the method used, which should facilitate coding of the computation.

Suppose there are $K$ strains that are shared and have a common ASV set. Let $r$ be the read count of the ASVs set associated uniquely with these strains. Let $n_j$ be the number of operons of strain $j$. Suppose the common cellular abundance is $\alpha$. The cellular abundances $c_j$ are given by $c_j = r_j/n_j$, where $r_j$ is the read count assigned to strain$j$. Then $c_1 = c_2 = \cdots = r_K/n_K = \alpha$, $\sum r_j = r$. Noting that $r_j = \alpha n_j$ we have $\alpha \sum n_j = r$ and hence $c_j = \alpha = r/\sum n_j$, $j = 1, \cdots, K$. Thus the proportion of ASV set counts for strain $k$ is $rn_k/\sum_{j=1}^{K} n_j$. This reduces to $r/K$ if all strains concerned have the same number of operons.

#### 5 Primers used for 16S rRNA gene and 23S rRNA gene identification and extraction

IUPAC codes are used to characterise primers that have variants in the base at particular locations. Most hypervariable regions have multiple variants. The 16S rRNA gene boundaries are not usually coincident between successive hypervariable regions, however the 23S rRNA gene Z regions were deliberately coincident and hence only the 3' boundary is specified for each Z region.

Table 21 presents results from unpublished original work by the second author. Multiple sets of quasi-randomly chosen 23S rRNA gene sequences from Refseq were created. Multiple sequence alignment was performed on each set and then per base Shannon entropy plots generated. Based on these, candidate locations for primers (low entropy regions of approximately 15 bases) were selected and the sequences in these regions examined to create a potential primer. Primers were then evaluated across separate random sets of sequences and progressively refined, resulting in the primers documented here. Full details are available in the thesis [4].

It will be noted that Table 21 has two entries for the 23S rRNA gene 3' boundary which are rather distant from each other. In this work the U2482R primer [5] has been used.

Table 20: Primers used for boundary identification for 16S rRNA genes

| Type | ID | Sequence | Comment |
|---|---|---|---|
| 16S rRNA gene_5' | 27F | AGAGTTTGATCMTGGCTCAG | |
| V2_5' | 97F_V | GGCGVACGGGTGAGTAA | |
| V2_3' | 338R | TGCTGCCTCCCGTAGGAGT | |
| V3_5' | V3F | CCAGACTCCTACGGGAGGCAG | |
| V3_3' | V3R | CGTATTACCGCGGCTGCTG | |
| V4_5' | 515F | GTGYCAGCMGCCGCGGTAA | Caporaso etal |
| V4_3' | 806R | GGACTACHVGGGTWTCTAAT | Caporaso etal |
| V5_5' | 784F | AGGATTAGATACCCT | |
| V5_3' | 908F_KW | ACTCAAAKGAATWGACGG | probeBase |
| V5_3' | 908R_M | TACGGYTACCTTGTTACGACTT | probeBase |
| V6_5' | V6F | TCGATGCAACGCGAAGAA | Chakravorty et al. |
| V6_3' | V6R | ACATTTCACAACACGAGCTGACGA | Chakravorty et al. |
| V7_5' | 1100.F16 | CAACGAGCGCAACCCT | |
| V7_3' | 1237F | GGGCTACACACGYGCWAC | |
| V9_5' | 1381R | GACGGGCGGTGTGTRCA | |
| V9_3' | 1492R | TACGGYTACCTTGTTACGACTT | |
| 16S rRNA gene_3' | 1492R | TACGGYTACCTTGTTACGACTT | |

Table 21: Primers used for boundary identification for 23S rRNA genes

| Type | ID | Sequence |
|---|---|---|
| 23S rRNA gene_5' | 129F_YV | CYGAATGGGGVAACC |
| Z0_3' | Z0F_84 | GAABTGAAACATCTHAGTA |
| Z1_3' | Z1F_136 | AGTAGYGGCGAGCGAA |
| Z2_3' | Z2F_384 | AGTACYGTGARGGAA |
| Z3_3' | Z3F_728 | WRATAGCTSGTWCTC |
| Z4_3' | Z4F_986 | GTTRGCTYRGAAGCAGC |
| Z5_3' | Z5F_1588 | AGGAAYTMKGCAA |
| Z6_3' | Z6F_1723 | TGAYRCCTGCCCRGTGC |
| Z7_3' | Z7F_1821 | TCCTAAGGTAGCGAAATTCCTTG |
| Z8_3' | Z8F_2144 | ACTGGGGYGGTYKCCTCC |
| 23S rRNA gene_3' | 2241R_H | ACCGCCCCAGTHAAACT |
| 23S rRNA gene_3' | U2482R_RM | CCRAMCTGTCTCACGACG |
